# Structure and genomic organization of the human *DUX4* homologue bovine *DUXC*

**DOI:** 10.64898/2026.03.05.709741

**Authors:** Barış Yaşar, Tõnis Org, Marilin Ivask, Gamze Yazgeldi Gunaydin, Nina Boskovic, Ülle Jaakma, Juha Kere, Ants Kurg, Shintaro Katayama

## Abstract

**Background:** *DUXC* is a multi-copy transcription factor gene found within a long tandem repeat locus in several Laurasiatherians. It is suggested to be functionally similar to human DUX4 because of its shared C-terminal domain and its close phylogenetic relationship to DUX4. *DUX* family genes are transiently expressed in preimplantation embryos of placental mammals. However, early embryo-derived cDNA proof for *DUXC*, which is needed for its further functional characterization, has not been reported so far.

**Results:** Our study provides a full-length sequence of *DUXC* mRNA, derived from the 8-cell stage in vitro fertilization (IVF) bovine embryos, containing double homeobox and 9aa transactivation domain (9aaTAD)-encoding sequences. Identified *DUXC* sequence uncovered a first exon that was not previously annotated. We showed that *DUXC* mRNA levels are independent of the embryonic transcription at the 2-, 4-, and 8-cell stage, whereas its decline, observed from the 8-cell stage onwards, is minor embryonic genome activation (EGA)-dependent. We also investigated the genomic organisation of the *DUXC* array in eight different cattle breed assemblies, revealing polymorphic internal repeats flanked by an incomplete distal unit at the telomeric end and a much shorter unit at the proximal end of the *DUXC* array. Despite the presence of a putative polyadenylation signal downstream the distal unit, we presented evidence for the expression of internal but not distal *DUXC* in early bovine IVF embryos.

**Conclusions:** *DUXC* is a potential bovine EGA inducer, supported by its expression at peak levels at pre-EGA stages followed by a decrease with a dependency on minor EGA.

## Background

*Double homeobox* (*DUX*) family genes belong to the PRD-like (PRDL) class transcription factor (TF) [1] and can be categorized into two groups based on the presence of an intron. These groups include intron-containing *DUXA, DUXB, DUXC*, and *DUXB-like* (*Duxbl*), and intron-lacking *Dux* and *DUX4* [2]. *DUXA* and *DUXB* were identified as the first intron-containing genes [3]. This was followed by the identification of *DUXC* and *Duxbl*, the former of which was initially identified in bovine (*Bos taurus*), dog (*Canis familiaris*), and armadillo (*Dasypus novemcinctus*) [2]. An earlier model proposed that *DUX4* and *DUXC* co-existed in a common ancestor [2] but later analyses supported a model in which an ancestral *DUXC* was retained in the genomes of Laurasiatheria, including bovines, and gave rise to *DUX4* retained in primate lineages [4]. It is hypothesized that retrotransposition and amplification of *DUXC* led to *DUX4* [2, 5], which was identified and named [6] after uncovering its tandemly repeated genomic organization through sequencing of the D4Z4 array [7]. Similar to *DUX4, DUXC* was also reported to be organized in tandem repeats [5], marking it as the only intron-containing *DUX* family gene located in a repeat array. Unlike *DUXA, DUXB*, and *Duxbl, DUXC* encodes a C-terminal domain that is conserved with *DUX4* and *Dux* [2, 5]. Moreover, homeodomains of DUX4 and DUXC are phylogenetically the most closely clustered among all DUX proteins [5]. Because *DUXC* does not co-occur with *Dux* or *DUX4* in the genome [5] and is closely related to the latter, DUXC and DUX4 are likely to be molecular counterparts of each other in different species [4].

DUX4 was identified as the key TF for regulating embryonic genome activation (EGA) in human in vitro fertilization (IVF) embryos [8, 9], with the mRNA induction observed at zygote stage [10, 11], and the enrichment of its binding motif identified in the promoters of EGA genes [8, 12]. Despite being suggested as a potential critical factor also for bovine EGA [13], *DUXC* gene structure has only been predicted based on similarity of the protein-coding sequences [2, 4, 5]. Current gene predictions for *DUXC* lack a gene symbol in both Ensembl [14] and NCBI [15] gene databases. Similarly, no TF binding motif exists for DUXC in a database [16] that stores high-quality DNA binding profiles. Providing an experimentally validated gene structure for *DUXC* is necessary for future functional characterization, for example to define its DNA-binding properties and contribute to TF motif collections. Moreover, computational gene predictions performed through genome-wide annotation pipelines are error-prone. Consortia, such as FANTOM [17] and H-Inv [18], demonstrated that full-length cDNA clones contain precise 5’ and 3’ end information, and help identify novel isoforms, revealing the actual structure of expressed genes. Therefore, sequence-level characterization of *DUXC* mRNA through molecular cloning is a prerequisite for elucidating its function during bovine EGA.

Although RNA-seq is a common method nowadays, there is a pitfall in investigating the structure and expression of *DUXC*. Because *DUXC* is a multi-copy gene, aligning RNA-seq reads to its locus often results in multi-mapped reads. Mappers use different approaches for aligning reads to multi-copy genes. For reads that map equally well to multiple genomic locations, mappers can randomly select one alignment [19], or they can consider them unmapped if the number of multiple alignments exceeds a certain threshold [20]. Computational tools that count reads might also ignore the multi-mapped reads [21], resulting in signal loss from such genes. For this reason, accurate quantification of *DUXC* transcripts in early embryos requires a *DUXC* locus-focused quantification approach.

*DUX* family genes are located in telomeric and pericentromeric regions of the chromosome [22]. Telomeric regions consist of repeats and high GC content, making short-read sequencing inefficient for studying such genomic regions [23]. On the other hand, long-read sequencing provides enhanced resolution of genomic regions with repetitive DNA sequences. While ultra-long reads made it initially possible to sequence several D4Z4 repeats [24], advancement in the sequencing technologies allowed complete resolution of the highly repetitive regions [25]. For this reason, studying a locus or gene found within a repeat array requires the use of long-read sequencing-based genome assemblies.

Here, we provide cDNA-level evidence for the expression of *DUXC* in the 8-cell bovine IVF embryos and update the structure of *DUXC*, revealing a previously unexplored first exon upstream of predicted gene models. We uncover the genomic organization of *DUXC*, arranged in tandemly repeated units in several cattle breeds, where the identified first exon was conserved. We also show that *DUXC* mRNAs detected in the 2- and 8-cell bovine IVF embryos are of internal but not distal unit origin. Our analysis demonstrates expressed *DUXC* mRNAs at the 2-, 4-, and 8-cell stage to be independent of transcription inhibition. In contrast, we show that the reduction of *DUXC* transcript levels at the 8-cell stage is dependent on minor-EGA. Collectively, these results improve the characterization of *DUXC* by illustrating its conserved genomic organization across breeds with an accurate gene structure, elucidating the repeat origin for its expression, and providing further insight into its relationship with EGA.

## Results

### Human DUX4 homologue bovine DUXC putatively binds to the promoters of genes activated during bovine EGA

In our recent work, we discovered the de novo motif 5’-TAAYCYAATCA-3’ that was enriched in the promoters of transcripts exclusively upregulated during bovine EGA [26]. Focusing on genes and proteins annotated in the bovine genome, we highlighted DUXA and PHOX2A as potential binders of this motif, identifying them as putative inducers of bovine EGA. Notably, this de novo motif also showed strong similarity to the known binding motif of proteins from other species, one of which is human DUX4 (MA0468.1; E-value = 6.1×10^−6^; q-value = 1.1×10^−5^; Fig. 1A; Supplementary Table 1).

**Figure 1.**
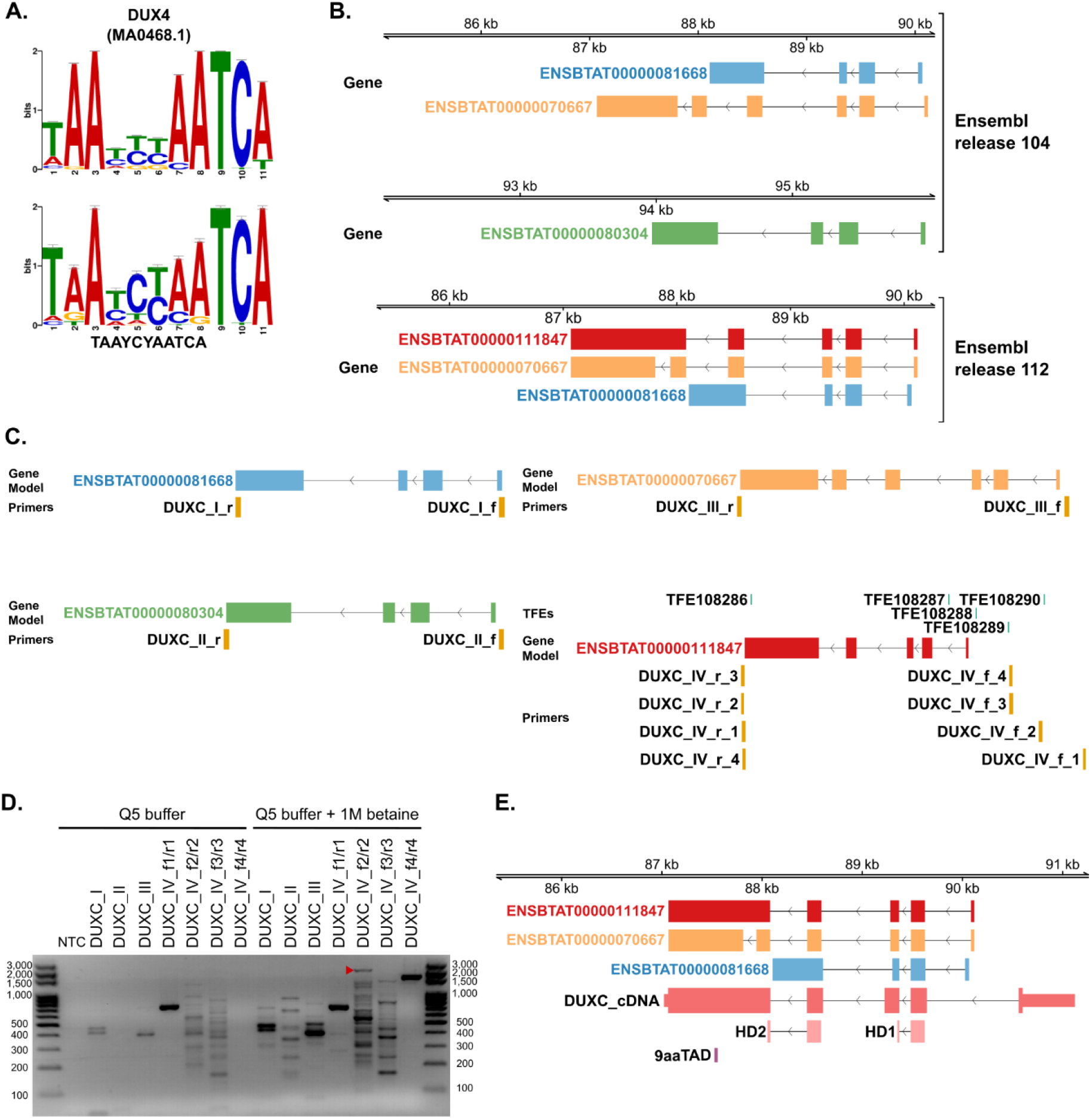
Molecular cDNA cloning of *DUXC* reveals a previously unannotated upstream exon. **(A)** De novo motif found in the promoters of 16-cell-specific genes upregulated following bovine EGA is shown at the bottom, with human DUX4 binding motif shown above it. **(B)** *DUXC* gene models from Ensembl release 104 and 112. **(C)** Gene models and their corresponding primer pairs used for PCR reactions. TFEs represent the transcript far 5’-ends detected in the *DUXC* locus based on the reanalysis of the dataset generated in our recent study [26] **(D)** Agarose gel showing the PCR-amplified *DUXC* fragment marked by an arrowhead. PCR reactions were performed using the primers in the absence (left) or presence (right) of 1M betaine. Lanes were labelled with the gene model targeted for amplification together with its primer pair. **(E)** Comparison of the *DUXC* gene models from Ensembl release 112 (in blue) with our cDNA cloning-verified *DUXC* (in coral red). Homeobox-encoding homeodomains 1 and 2 were labelled as HD1 and HD2 (in light pink), respectively. DNA sequence coding for the 9aaTAD was marked in light magenta.

Although DUX4 is not encoded in the bovine genome, earlier findings [2, 4, 5] suggesting the presence of its homologue, DUXC, prompted us to speculate that this consensus sequence is likely bound by DUXC. Despite being a putative bovine EGA inducer based on our motif analysis (Fig. 1A), proper annotation of *DUXC* is missing as only computational gene predictions exist in Ensembl [14]; e.g. ENSBTAT00000111847.1 transcript in an unnamed gene was predicted from human DUX4L2 (UniProt/SWISSPROT: P0CJ85) without any transcriptional evidences in bovine. For this reason, we set out to perform molecular cloning of *DUXC* cDNA derived from the 8-cell stage bovine IVF embryos in order to provide experimental evidence for the full-length *DUXC* cDNA sequence.

### Isolation of *DUXC* mRNA from early bovine embryos reveals a novel upstream exon

Since the gene symbol for *DUXC* is missing in the Ensembl gene annotation, we performed tblastn search using the putative protein sequence of porcine DUXC [27] to determine the genomic location of bovine *DUXC*. At the pericentromeric locus on chromosome 7, the UCSC Genome Browser showed three distinct Ensembl predictions (release 104), two of which appeared as tandemly repeated copies with identical structure (Supplementary Fig. 1A).

We selected non-redundant transcript predictions (ENSBTAT00000081668, ENSBTAT00000070667, ENSBTAT00000080304) from Ensembl release 104 but also included a distinct prediction (ENSBTAT00000111847) from the release 112 to design PCR primers (Fig. 1B). We initially designed PCR primers for gene models from Ensembl release 104 (Fig. 1C). After reanalyzing our recently generated RNA-seq dataset [26] with a TFE-based (transcript far 5’-ends) approach [12, 28], which can discover ultimate 5’ ends of transcripts, we found TFEs in the upstream of the transcript model ENSBTAT00000111847 (Fig 1C, lower right). Therefore, we also designed primers considering these TFEs to test the transcription initiation from a potential novel exon located upstream of the gene models (Supplementary Table 2). The expected PCR product lengths were 733 nucleotides (nt) for DUXC_I, 813 nt for DUXC_II and 1382 nt for DUXC_III.

For DUXC_IV, using primer pairs f1/r1 through f4/r4, the expected lengths were 2620 nt, 2067 nt, 1670 nt and 1622 nt, respectively. Our initial attempts to perform PCR for *DUXC* with Phusion polymerase did not result in amplification of the target fragment, which was also observed for a template DNA with a moderate GC content [29], similar to that of *DUXC* (Supplementary Table 3). As a result, we opted for Q5 polymerase, as it was proven to amplify a gene with GC content similar to *DUXC* [29]. Among the primer pairs tested (DUXC_I-IV), DUXC_IV (primer set f2/r2) produced a fragment at the expected length in the presence, but not in the absence of betaine (Fig. 1D; Supplementary Fig. 1B). Notably, even though DUXC_I showed a band around its expected length, Sanger sequencing of this fragment after cloning showed that this was a nonspecific band. Molecular cloning of the fragment DUXC_IV amplified by the primer set f2/r2 revealed a *DUXC* structure distinct from its in silico predictions, with the first exon located further upstream than in other *DUXC* models (Fig. 1E). Furthermore, we verified the presence of complete two homeobox and the sequence encoding the nine amino acid transactivation domain (9aaTAD) [30] in the cloned *DUXC* sequence, suggesting that it is functional as a double-homeodomain transcription factor (Fig. 1E). In summary, we provide a more reliable structure, including the UTRs, of *DUXC*, validated by cDNA cloning, which will facilitate further characterization of its role during bovine EGA.

### Minor EGA-blocker prevents a decline in *DUXC* transcript levels at the 8-cell stage

Having a more reliable, cDNA-cloning confirmed annotation of *DUXC*, we set out to reanalyze publicly available RNA-seq datasets to examine the dynamic expression of *DUXC* across various developmental stages. First, we utilized a dataset [31] composed of stages with normal IVF embryo development in addition to those subjected to certain treatments to produce transcriptionally blocked embryos (TBE), including embryos with minor EGA-block.

Our reanalysis showed increased levels of *DUXC* transcripts following fertilization at the zygote stage with a subsequent decline at the 2-cell stage (Fig. 2A). Interestingly, we detected increased expression at the 4-cell stage, after which decreased levels of *DUXC* mRNAs were observed at almost all subsequent developmental stages except for one inner cell mass (ICM) replicate (Fig. 2A). We also confirmed the mapping of RNA-seq reads to the *DUXC* first exon identified in this study (Supplementary Fig. 2; Fig. 1E). As references, we included other PRDL homeobox genes *ARGFX, DUXA, LEUTX*, and *TPRX*s that peak their expression during bovine EGA, and *NOBOX* that is primarily expressed in oocytes [26]. For RNA-seq reanalysis, we used cDNA-confirmed gene annotations for these genes that we previously reported [32]. Even though embryonically-expressed control PRDL homeobox genes were affected by the α-amanitin-mediated transcription inhibition at the 4- and 8-cell stage, *DUXC* transcript levels were not affected by this inhibition at any stage (Fig. 2A).

**Figure 2.**
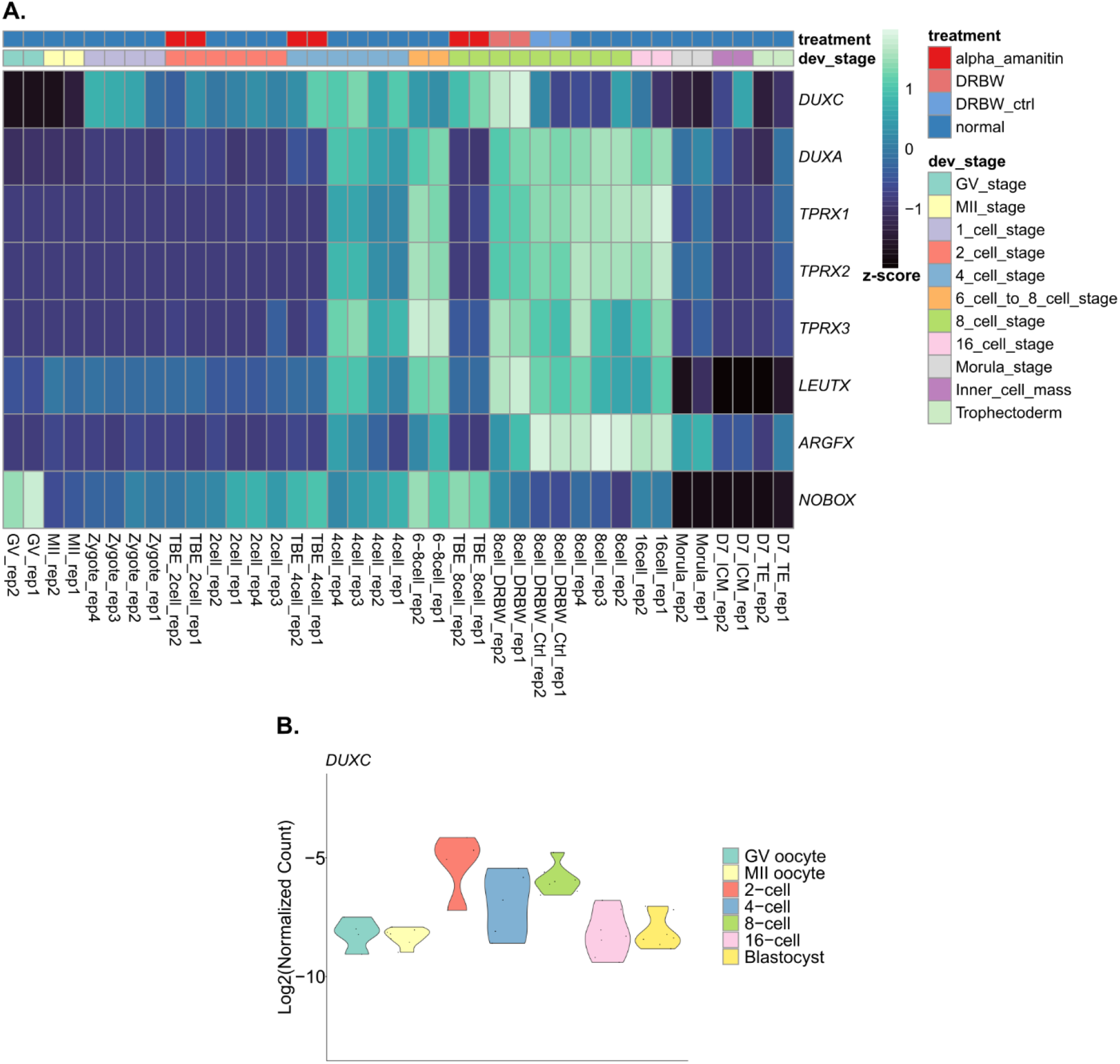
*DUXC* mRNA is observed at pre-EGA stages with its clearance depending on minor EGA-block. **(A)** Heatmap showing the expression of *DUXC* and other PRD-like (PRDL) homeobox genes cloned earlier [32] across bovine early development using the RNA-seq dataset generated in a recently published study [31]. Treatments include alpha_amanitin and DRBW. The former represents the α-amanitin treatment used to block embryonic transcription until the collection at the pre-EGA stages (2- and 4-cell stage) or at the EGA stage (8-cell stage). These embryos are also referred to as TBE or transcriptionally-blocked embryos. DRBW represents DRB wash, in which 5,6-dichlorobenzimidazole 1-β-D-ribofuranoside (DRB) was used to obtain the 8-cell stage embryos with minor EGA-block. DRBW_ctrl represents the control group for this treatment. **(B)** Violin plot showing the expression of *DUXC* based on spike-in RNA normalization using the RNA-seq dataset from our recent work [26].

Inhibition of minor EGA (denoted by DRBW) caused an increase in the levels of *DUXC* mRNAs at the 8-cell stage, where *DUXC* transcript levels start to decrease in normal IVF embryo development in this dataset. A slight increased expression of *LEUTX* and *TPRX3* was also observed in minor EGA-blocked embryos collected at the 8-cell stage as compared to the corresponding control embryos. In contrast, minor EGA inhibition caused a reduction in *ARGFX* expression.

Next, in order to provide a more accurate expression profile of *DUXC* during bovine IVF embryo development, we used the dataset from our recent study [26], where we employed a *DUXC*-centric reanalysis approach, masking all except for one copy of *DUXC*. Even though this dataset did not include the zygote stage, we observed the highest expression at the 2-cell stage with a slight decrease at the 4-cell stage (Fig. 2B), unlike in the previous dataset whose normalization does not utilize spike-in RNAs (Fig. 2A). The levels of *DUXC* dropped noticeably at the 16-cell stage with similar levels being observed at the blastocyst stage. We also observed RNA-seq reads of *DUXC* origin in oocytes, as judged by our previously defined cutoff of -10 to indicate expression in the violin plot for this dataset [26] (Fig. 2B). Collectively, we show the highest levels of *DUXC* mRNA to be at pre-EGA stages with its levels not being dependent on the embryonic expression and confirm the transcription from the novel first exon.

### Bovine *DUXC* is organized in tandem repeat units flanked by incomplete repeat units

In order to characterize the array *DUXC* is found in, we investigated this locus using both Hereford and Holstein genome assemblies, revealing the organization and number of tandem repeat units (Fig. 3). We used an independent Holstein bovine assembly [33] (named NCBA_BosT1.0) based on long-read sequencing. In addition, we also investigated the reference Hereford bovine assembly (ARS-UCD2.0). We performed in silico PCR [34] and showed that there are six and 18 complete repeat units in Hereford and Holstein, respectively (Fig. 3A; Supplementary Table 4). The length of each unit slightly varied in Holstein cattle, with some deletions of up to 400 bp observed in some units (Fig. 3A). Interestingly, the median of the repeat unit was slightly lower in Hereford cattle (Fig. 3A).

**Figure 3.**
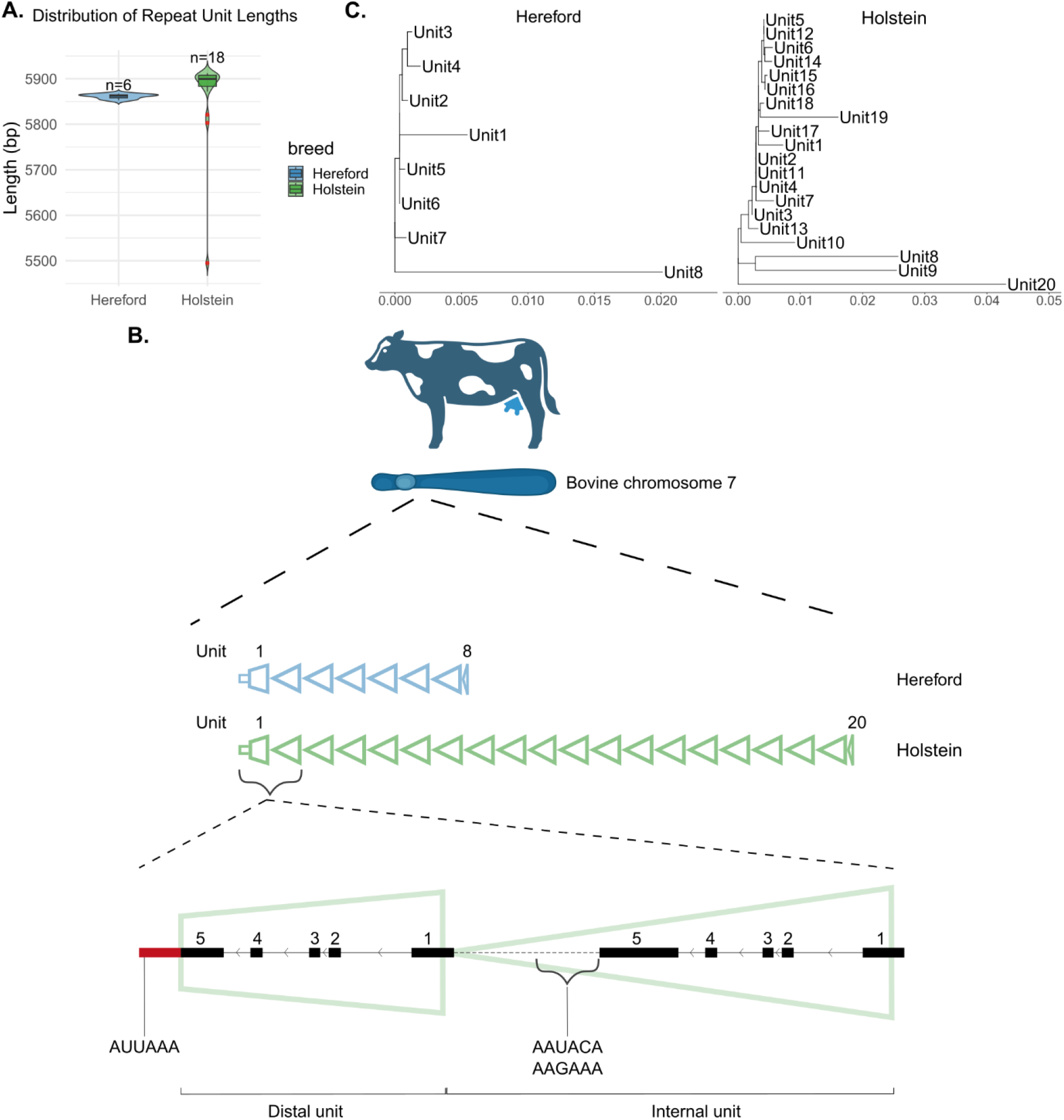
Investigation of *DUXC* loci in Hereford and Holstein cattle uncovers tandemly repeat organization of *DUXC*. **(A)** Distribution of internal repeat unit lengths in each breed. **(B)** Schematic overview of the *DUXC* genomic locus in chromosome 7. Each triangle represents one repeat unit with the incomplete repeat units at both ends drawn as truncated triangles. First and last repeat unit numbers were marked. A zoomed in view of the distal and the immediate internal unit were shown to highlight the *DUXC* gene structure in each unit. *DUXC* is located on the negative strand. Each exon was marked with its corresponding exon number. The variant portion of the exon 5 in the distal unit was coloured in red. The putative polyadenylation signals present in the genomic sequence were also marked. **(C)** Phylogenetic trees representing the sequence similarity among *DUXC* repeat units within each breed.

Upon manual inspection of the locus, we observed an incomplete distal unit at the telomere end of the array in both cattle breeds, which we referred to as Unit 1 (Supplementary Text 1). This unit was not amplified by in silico PCR as it did not retain the sequence for the forward primer to anneal. The opposite end of the array in both breeds also contained incomplete and much shorter repeat units, neither of which were amplified by in silico PCR because of lacking the sequence for the reverse primer to anneal (Supplementary Text 2).

Based on these results, we visualized the repeat units together with our cDNA cloning-based *DUXC* structure to demonstrate the genomic organization of this locus (Fig. 3B). The distal *DUXC* located on chromosome 7 toward the telomeric end had a variant 3’ end, causing the downstream portion of the putative *DUXC* exon sequence to be altered (Fig. 3B, shown in red). In this variant portion of the sequence in both Hereford and Holstein genomes, we detected the putative polyadenylation signal AUUAAA (Fig. 3B; Supplementary Fig. 3), a common hexamer found in the 3’-UTRs of transcripts [35]. We also identified less frequent polyadenylation signals [35] such as AAUACA and AAGAAA in the downstream regions of the internal *DUXC* copies (Fig. 3B). These polyadenylation signals were also present downstream of the 3’-UTRs of putative internal *DUXC* copies in both Hereford and Holstein (Supplementary Fig. 3).

For each breed, we performed multiple sequence alignment using all units and built a tree to examine how they are related to each other (Fig. 3C). Our analysis showed that Hereford internal units (Unit 2-6) were closely similar but not identical as evidenced by very short branch lengths, whereas the incomplete units (Unit 1 and Unit 8) had much longer branch lengths (Fig. 3C, left). In Holstein, the most divergent unit was the one at the non-telomeric end of the array (Unit 20) with a long branch, while Unit 8 and 9 formed an intermediate subcluster (Fig. 3C, right). Overall, using in silico PCR we uncover the genomic organization of tandem repeat units at the *DUXC* loci.

### Internal but not distal *DUXC* is expressed in 2-cell stage IVF embryos

In order to determine whether the *DUXC* transcripts found in bovine early embryos are derived from the distal or internal unit, we extracted each genomic region, to which we aligned reads from a publicly available RNA-seq dataset [36]. Because we wanted to elucidate whether the deviated end portion of the distal *DUXC* (Fig. 3B) is found in expressed transcripts, we opted for a dataset [36] prepared using a method ensuring sufficient coverage of the 3’ ends [37].

As we noticed an obvious difference at the 3’-end of exon 5 between putative distal and internal *DUXC*, we used it as an indicator of which copy is expressed (Fig. 3B; Supplementary Fig. 4). Furthermore, we detected 51 bp deletion in Holstein distal *DUXC*, a region spanning the beginning of exon 5 and the preceding intron, making the distal and internal copies of *DUXC* distinguishable (Fig. 4A). We detected reads spanning exon 4 and 5 in internal but not in distal *DUXC*, suggesting the expression of internal *DUXC* in Holstein 2-cell embryos (Fig. 4B). In addition, we detected reads aligning to the part of exon 5 in internal *DUXC* whose sequence differed in the distal copy (Fig. 4B, Supplementary Fig. 4). Some portions of these reads aligned to the sequence shared between internal and distal *DUXC* copies (to the right of the vertical dashed line), while the remaining portions aligned to the internal *DUXC*-specific sequence (to left of the vertical dashed line), providing an additional evidence for the expression of internal *DUXC* copy (Fig. 4B). Furthermore, using a different dataset [31], we also observed mapped RNA-seq reads at the 3’ end of the internal *DUXC* at the 4-cell stage (Supplementary Fig. 2).

**Figure 4.**
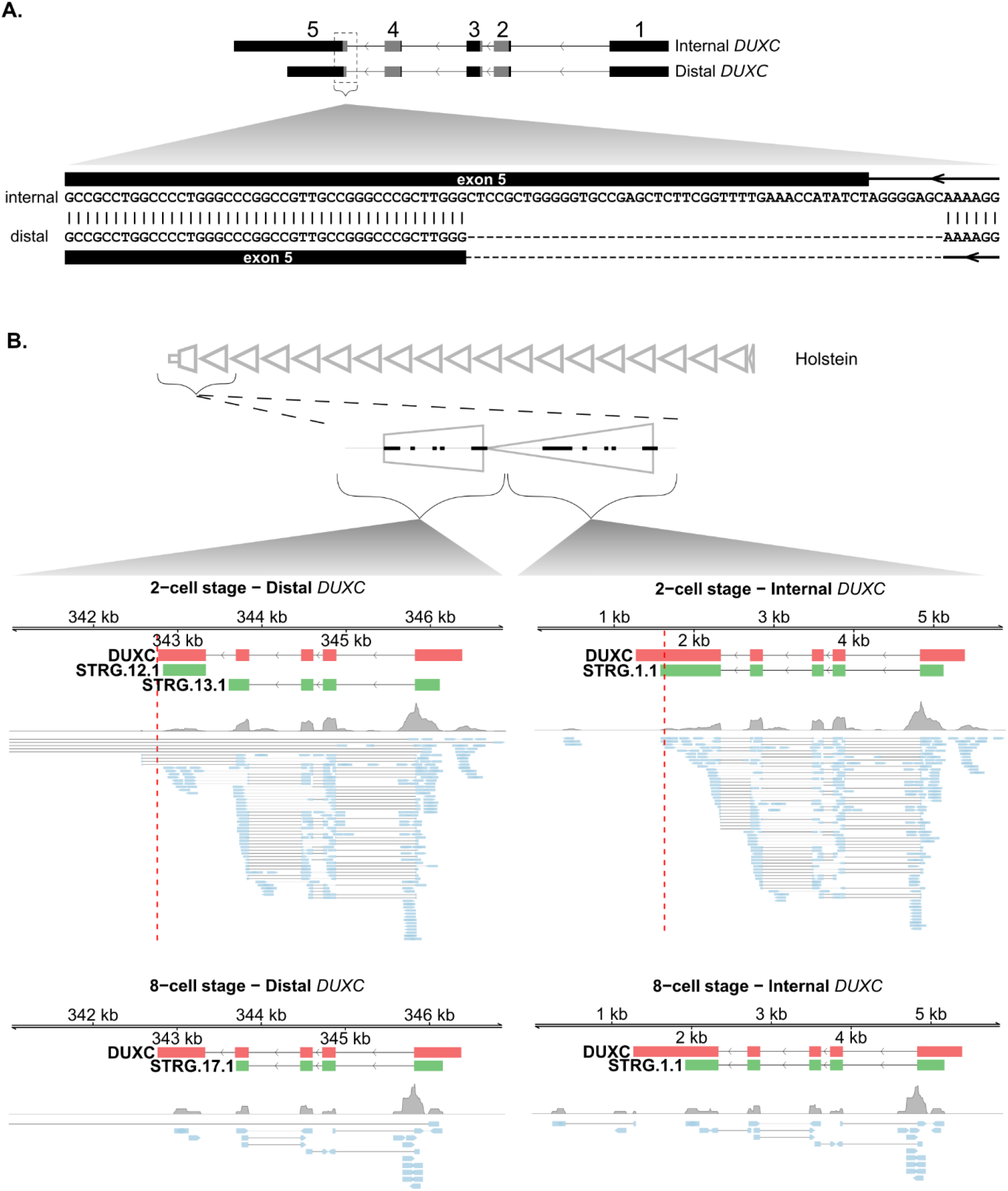
*DUXC* mRNA in bovine early embryos are derived from internal but not distal units. **(A)** Pairwise alignment of *DUXC* exon-intron junction at the distal unit (Unit 1) and internal unit (Unit 2). A portion of exon 5 and the preceding intron of Holstein cattle is shown. Continuous and dashed horizontal lines denote introns and the 51 bp deletion, respectively. The DNA sequences encoding homeodomain are coloured in grey. **(B)** Alignment of RNA-seq reads from a publicly available dataset [36] to the genomic region where the distal and internal *DUXC* gene annotation is found. RNA-seq reads were derived from the 2-cell (top) and 8-cell (bottom) bovine IVF embryos. The transcripts with STRG prefixes were annotated by StringTie [38] following mapping of RNA-seq reads to the shown loci. *DUXC* is located on the reverse strand. Annotation of *DUXC* was performed by the gmap [39] alignment of *DUXC* cDNA sequence to the shown loci. The red dashed vertical line indicates the region after which the sequence (to the left) starts to differ between putative distal and internal *DUXC. DUXC* cDNA (in coral red), assembled transcripts (green), read coverage (in gray) and RNA-seq reads (in blue) were shown at each unit.

In order to rule out the possibility that the distal *DUXC* might be expressed but with an alternative exon 5 located further downstream, we searched for reads that are split between exon 4 and a downstream region. For this purpose, we first ensured that Holstein genome assembly (NCBA_BosT1.0) had an assembled telomere at the end of chromosome 7, where *DUXC* is located, using telomere identification toolkit (tidk) [40] (Supplementary Fig. 5). Having extracted the genomic region of chromosome 7 spanning from the first nucleotide (telomeric end) up to the distal *DUXC*, we aligned RNA-seq reads to this region, allowing an intronic length equivalent to this distance as the extracted genomic region is shorter than the default intron length of HISAT2 [19] (Fig. 4B, left; Supplementary Table 5). However, we did not find any reads indicative of distal *DUXC* expression. Even though we identified four reads split between exon 4 and a downstream location of the distal region, it is unlikely to represent an alternative exon for distal *DUXC* (Fig. 4B, Supplementary Fig. 6A). These reads were likely derived from the internal *DUXC* copy as we observed the reads with the same IDs aligning to the internal unit split between exon 4 and 5 (Supplementary Fig. 6B).

We annotated *DUXC* by aligning our cDNA clone to each extracted locus, which resulted in a shorter last exon in the distal unit, in line with its diverged sequence at the 3’ end (Fig. 4B, left; Supplementary Fig. 4). Transcript assembly yielded the same intron-exon structure as our cDNA clone in both 2- and 8-cell embryos (Fig. 4B, right), providing an in silico confirmation of the gene structure we identified. In the 8-cell stage embryos, we observed a read that spanned exon 4 and 5, indicating that the *DUXC* transcripts observed at this stage were also derived from an internal unit or units (Fig. 4B). Only a small number of reads were present for other stages except for the blastocyst stage, which showed expression of regions overlapping with the exons and introns of *DUXC*, suggesting the expression of another feature (Supplementary Fig. 7). Consequently, we present evidence for the transcription of internal but not distal *DUXC* copy.

### Sequence identity and organization of *DUXC* arrays across eight cattle breeds are highly conserved

We extended our search for *DUXC* locus to several bovine breeds, utilizing long-read sequencing-derived genome assemblies, to assess the conservation and divergence of this locus (Supplementary Table 6). We used one of the internal Holstein *DUXC* repeat units (Unit 2) to define the boundaries of the array and extracted the *DUXC* locus from chromosome 7 of each breed (Supplementary Text 3; Supplementary Table 5). Following the in silico isolation of the array, we compared Jersey [41], Hanwoo [41], Holstein [33], Hereford, Tibetan [42], Mongolian, Tuli, and Wagyu [43], revealing the identity shared between the locus and the one immediately below it (Fig. 5A). Notably, we did not detect any *DUXC* repeat unit hits in Yunling [41], and nor any alignments between Yunling and any other breed.

**Figure 5.**
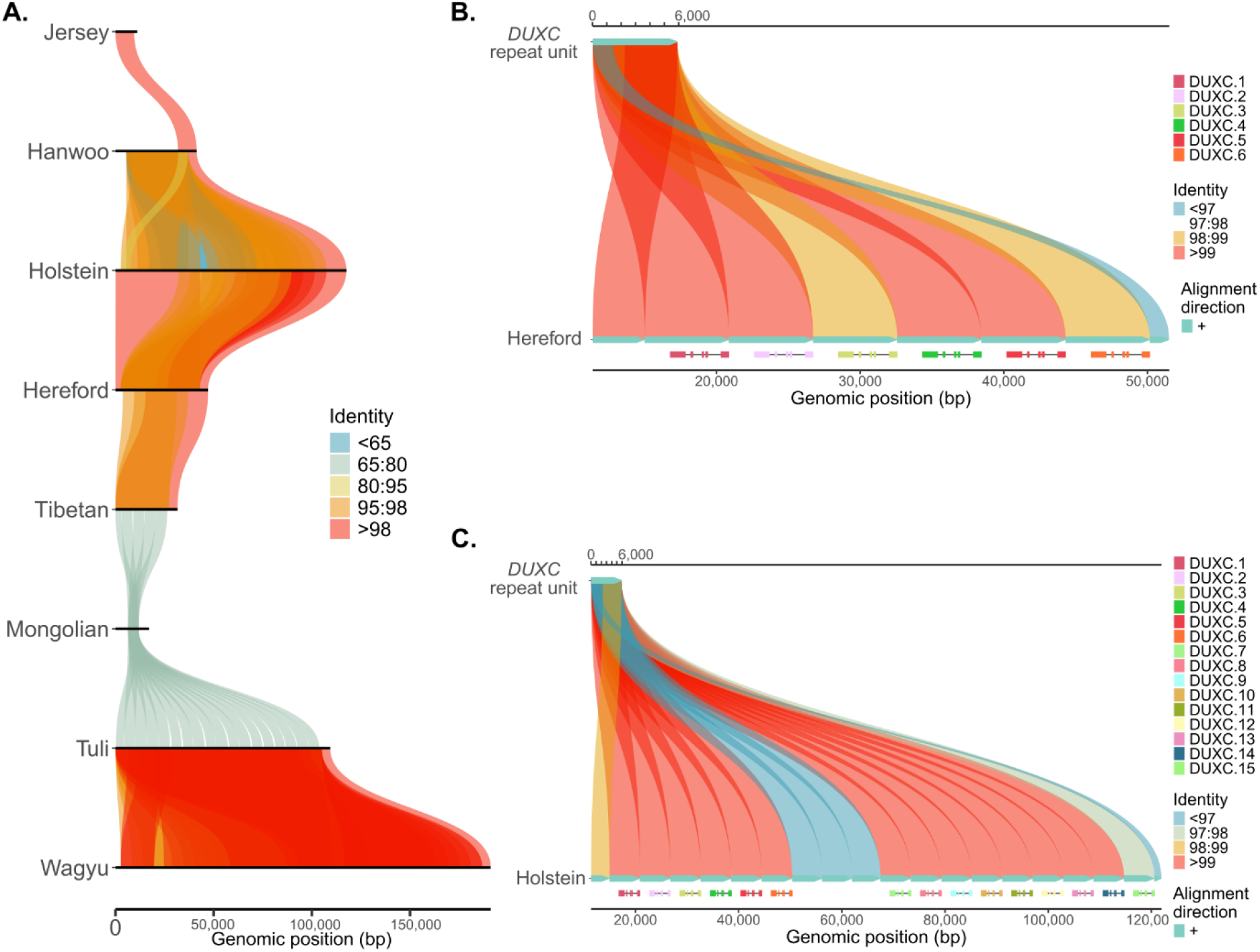
Comparison of *DUXC* locus in eight cattle breeds illustrates high sequence identity. **(A)** A comparative alignment of Jersey, Hanwoo, Holstein, Hereford, Tibetan, Mongolian, Tuli, and Wagyu visualized in subsequent order. Each locus is compared with the locus beneath it as stacked alignments [44]. **(B-C)** Holstein internal *DUXC* repeat unit (Unit 2) alignment against the entire *DUXC* locus of **(B)** Hereford **(C)** and Holstein cattle.

Following the extraction of *DUXC* locus, we examined its composition in each breed by aligning the same repeat unit (Unit 2) that defined the array boundaries, this time to identify each repeat unit within the extracted array (Supplementary Table 7). We observed the smallest *DUXC* array in Jersey, consisting of one repeat unit with a truncated downstream part and a small portion of the next repeat unit, together providing a sequence sufficient for *DUXC* annotation (Fig. 5A, Supplementary Fig. 8A). Almost the entire array in Jersey showed very high identity to Hanwoo, with more than 98% sequence identity. Hanwoo *DUXC* locus revealed five complete repeat units each with an intact *DUXC* gene structure and incomplete repeat units at each end of the array (Supplementary Fig. 8B). In the comparison between Hanwoo and Holstein, the identity dropped to below 65% in some parts of locus, whereas other regions showed 80-90%, 95-98% or even higher identity (Fig. 5A).

As having the second largest *DUXC* loci among all breeds investigated, Holstein contained 18 complete repeats but lacked a gene annotation in three of the internal repeat units, where the sequence identity was slightly lower than other internal repeat units (Fig. 5B). Holstein *DUXC* array was approximately 2.5 times longer than that of Hereford, which shared a sequence identity of more than 80% with both Holstein and Tibetan, reaching up to 98% in some regions (Fig. 5A). Complete *DUXC* repeat units from Holstein showed a minimum of 98% match, revealing the conservation of the internal units (Fig. 5C).

The Mongolian genome assembly contained the smallest *DUXC* array after Jersey, and showed nucleotide conservation in the range of 65% to 80% with Tibetan and Tuli genome assemblies (Fig. 5A). The beginning of the extracted locus coincided with the first nucleotide of chromosome 7 in the Jersey, Tuli, and Tibetan assemblies (Supplementary Table 5). Our approach of annotating *DUXC* using the Holstein-derived *DUXC* cDNA sequence did not produce a gene structure in the Mongolian *DUXC* locus, probably due to sequence divergence or partial assembly of the *DUXC* locus in this breed (Supplementary Fig. 8C).

The array in Tuli genome assembly had uninterrupted, complete internal repeat units in consecutive order, each with an annotated *DUXC* (Supplementary Fig. 8D). Unlike other breeds, the distal unit in Tuli had an annotation for *DUXC* with the last exon being slightly shorter than others. Four complete internal repeat units with partial ones at the beginning and end of the array was observed in the Tibetan *DUXC* locus (Supplementary Fig. 8E).

Wagyu genome assembly was notable not only for having the longest *DUXC* array but also for having the greatest number of annotated *DUXC* genes, with 28 copies present (Supplementary Fig. 8F). Unlike loci in other breeds, Wagyu *DUXC* locus was interrupted by a region where no match was found for the *DUXC* repeat unit. We also confirmed the across-breed conservation of the *DUXC* first exon that we identified in this study (Supplementary Fig. 9; Fig. 1E). Collectively, we present the genomic structure of the *DUXC* locus and demonstrate the conservation and divergence of *DUXC* repeat units across multiple breeds.

## Discussion

In this study, we provide the first experimentally validated annotation of bovine *DUXC*, a double-homeobox transcription factor previously represented only by computational predictions of the protein coding region. By cloning full-length *DUXC* cDNA from 8-cell stage embryos, we also identified a novel upstream exon and confirmed the presence of two homeodomains and a functional transactivation domain, establishing its potential role in early embryonic genome activation (EGA). Our expression analyses revealed that *DUXC* transcripts peak at pre-EGA stages, remain unaffected by major transcriptional inhibition, and are modulated by minor EGA-blocking treatments, suggesting a unique regulatory mechanism distinct from other PRDL homeobox genes. Furthermore, we uncovered the genomic organization of canonical *DUXC* as tandem repeat arrays on chromosome 7, demonstrated that internal copies, but not distal variants, are transcriptionally active in early embryos, and showed that this locus is highly conserved across multiple cattle breeds despite variation in array size. Collectively, these findings refine the structural and functional understanding of *DUXC* and highlight its evolutionary conservation and potential importance in bovine early development.

Gene prediction as a genome annotation can be performed by evolutionary comparisons that consider conserved synteny, intron-exon junctions and functional protein domains [5] or by computational methods that utilize datasets reflecting expressed genes [45]. These insightful approaches must still be complemented by experimental evidence using biological material originating from the tissue or developmental stage being investigated to confirm existing or identify new transcription start sites (TSSs).

Identifying the TSS or very 5’ end of a gene helps better locate its promoter, enabling the study of regulatory elements found there. Previously, transcript far 5’-ends (TFEs) of genes expressed in early human embryos were identified [12]. Molecular cloning of several genes guided by TFEs revealed previously uncharacterized isoforms [12, 46]. For example, the *LEUTX* gene model confirmed by cDNA cloning activated a greater set of genes than the RefSeq prediction [47]. Similarly, through molecular cDNA cloning, we demonstrated expression of several PRDL homeobox gene isoforms in early bovine embryos [32].

Performing TFE analysis using the dataset generated recently [26], we identified a potential TSS for *DUXC* (Fig. 1C). Guided by these TFEs, we aimed to amplify *DUXC* cDNA. Because of the GC-rich sequence composition of *DUXC*, the PCR assay required optimization, including the selection of a DNA polymerase better suited for GC-rich templates [29], as well as the addition of an agent to minimize the secondary structure formation mediated by GC-rich sequences [48]. Following successful amplification of *DUXC* cDNA (Fig. 1D), molecular cloning revealed a novel first exon for *DUXC*. Our experimentally validated sequence provides a more reliable gene structure for *DUXC*, and contains sequences encoding its functional domains; a double homeodomain and 9aaTAD. Even though primer pair f1/r1 did not amplify DUXC_IV (Fig. 1C-D), it is likely to be caused by the GC-rich content of the sequence, and we cannot rule out the possibility of an exon 1 that extends further upstream, beyond the 5’ end of *DUXC* we validated experimentally.

The discovery of a functional *DUXC* gene with a novel upstream exon and its peak expression at pre-EGA stages suggests that this transcription factor may act as an early transcriptional regulator in bovine embryos. Its independence from major transcriptional inhibition implies that *DUXC* transcripts are maternally deposited or activated through mechanisms preceding canonical EGA, positioning DUXC as a potential initiator of developmental programs rather than a downstream effector. Cytoplasmic polyadenylation [49] is a possible activation mechanism, and its potential involvement in regulating *DUXC* transcripts should be investigated.

Investigation of bovine *DUXC* locus revealed an array composed of highly similar but distinct repeat units organized in tandem similar to the genomic organization of D4Z4 array in humans (Fig. 3) [23, 50, 51]. Interestingly, we observed a putative polyadenylation signal AUUAAA downstream the distal unit, prompting us to speculate a transcription activity from the distal unit (Fig. 3C). Expression of two distinct intron-containing *DUX4* isoforms extending into a polyadenylation signal AUUAAA, the same signal we observed in bovine, was discovered only in facioscapulohumeral muscular dystrophy (FSHD) samples [52]. In the disease state, this polyadenylation signal was suggested to help mRNA extend its half-life to enable its translation [52]. Even though this observation was only made in a human disease setting but not in an embryonic development setting in bovine, we examined whether functional distal *DUXC* could be expressed to potentially utilize AUUAAA signal in the same way it is expressed in FSHD samples. Our investigations did not show any evidence for the expression of distal *DUXC* (Fig. 4) but we observed less-frequent putative polyadenylation signals [35] in the downstream region of internal *DUXC* copies (Fig. 3C; Supplementary Fig. 3). Currently, RefSeq gene models for DUX4 originate from FSHD samples and early human embryo-derived full-length *DUX4* sequence is not yet reported.

Comparative analysis of *DUXC* locus across cattle breeds revealed to what extent the *DUXC* array is assembled. Our comparative analysis allowed us to demonstrate a similar genomic organization almost in all breeds, with internal repeat units flanked by incomplete ones at both ends of the array (Fig. 5; Supplementary Fig. 8). In line with the strength of long-read sequencing in resolving repeated sequences, assemblies with longer contig N50 usually had a longer assembled *DUXC* locus. For example, Holstein (75x; PacBio RSII only) and Wagyu (286.9x; PacBio HiFi and Nanopore ultra-long > 100kb reads) showed both the longest *DUXC* array and contig N50 length (Supplementary Table 6; Fig. 5A). Genome coverage is another factor influencing the assembly of a genomic region, as demonstrated by the Holstein breed, which had a higher genome coverage and consequently a longer *DUXC* array assembled, despite having almost the same contig N50 length as Tibetan cattle (Supplementary Table 6).

In this study, we report mapping a single internal Holstein *DUXC* repeat unit (Unit 2) to the entire *DUXC* locus from other breeds. Previously, predicted *DUXC* copy numbers were reported in four distinct cattle breeds that we did not investigate, reporting copy numbers much greater than our analysis showed [4]. This earlier study does not claim to report absolute copy numbers but to provide evidence for amplification of *DUXC*, demonstrating its multi-copy organization [4]. Even though we did not use the same cattle breeds, the discrepancy in copy numbers likely arises because of methodological differences, specifically the reliance on either short-read or long-read sequencing. *DUX4* located in the D4Z4 array was shown to exist in varying copy numbers in healthy human individuals [51]. It is likely but not shown yet that inter-individual *DUXC* copy number differences are also observed in bovine.

DUXC and DUX4 were suggested to be functional homologs [4] and to have very similar binding motifs [13]. Although its binding motif is not deposited in databases of DNA binding profiles, the observation that the de novo motif 5’-TAAYCYAATCA-3’ enriched in the promoter of genes activated during bovine EGA shows a close match to the DUX4 binding motif suggests DUXC as a potential binder of this motif (Fig. 1A). Crystal structure of the DUX4-DNA complex identified 5’-TAAT-3’ and 5’-TGAT-3’ as the core sequences first and second homeodomain (HD1 and HD2) bind in a head-to-head orientation, respectively [53]. The binding of HD1 and HD2 to DNA is strong when these core sequences are separated by three but not two nucleotides [53]. In our de novo motif 5’-TAAYCYAATCA-3’, we observed the same core sequences 5’-TAAT-3’ and 5’-TGAT-3’, separated by tree nucleotides, supporting DUXC DNA binding orientation similar to that of DUX4. On the contrary, while HD2 was shown to be conserved among its homologs in other species, HD1 was distinct in primates [53], raising the possibility that DUXC HD1 has a different sequence specificity. Further research is needed to define the precise sequence specificity of DUXC.

Our observation showing that the detected *DUXC* transcripts are not affected by transcription inhibition was made earlier for the 8-cell stage bovine embryos [54]. By reanalyzing a publicly available dataset [31], we confirmed this observation and further demonstrated that this is also true for the 2- and 4-cell stage bovine embryos (Fig. 2A). Furthermore, the inability of the 8-cell stage embryo to downregulate *DUXC* transcripts upon minor EGA inhibition suggests that this earlier wave of genome activation might be activating genes that are involved in clearance of *DUXC* transcripts. Therefore, *DUXC* mRNAs are likely needed within a short time window during early development. While these results show relative expression of *DUXC* at every stage of embryonic development, our recent dataset [26], although covering fewer developmental stages, presents a more accurate expression profile as it utilizes unique molecular identifiers (UMIs) [55] for correcting PCR duplicates and added spike-in RNAs for normalization [28] (Fig. 2B).

*DUXC* is thought to have retrotransposed to give rise to *DUX4* in humans and *Dux* in mice [2, 5]. For retrotransposition events to take place, retrotransposed genes must be expressed in germ cells, through which the resulting gene can be passed on to new generations and maintain its existence [3]. In line with this, we detected the expressed *DUXC* transcripts in oocytes at low levels (Fig. 2B). Because of the C-terminal domain that is conserved between DUX4 and DUXC, this domain was suggested to act as a transcriptional activator [4] and was later shown to recruit [56] transcriptional co-activators [57] p300/CBP. Consistently, the cloned *DUXC* cDNA was predicted code for 9aaTAD [30] in its 3’ end (Fig. 1E). Canine DUXC was also shown to activate 2-cell-like genes in mouse muscle cells, supporting the transcription activation capacity of DUXC [58]. Furthermore, evidence from overexpression experiments and open chromatin data [13] point to the role of DUXC during EGA. Collectively, DUXC is a PRDL homeodomain TF with potential transcription activation activity that might be involved in bovine EGA.

Although our study provides the first experimentally validated annotation of bovine *DUXC* and insights into its expression dynamics, several limitations should be acknowledged. First, the functional role of *DUXC* during early development remains inferred from structural features and expression patterns rather than direct functional assays; loss-of-function or gain-of-function experiments will be necessary to confirm its regulatory impact on EGA. Second, while we demonstrated transcription from internal repeat units, the possibility of distal copies being expressed under specific conditions cannot be fully excluded due to limited RNA-seq coverage. Finally, our comparative analysis across breeds was based on available genome assemblies, which vary in completeness and may underestimate structural diversity in the *DUXC* locus. Since each genome assembly has a different genome coverage and contig N50 values, we can only compare the given assemblies but cannot deduce the entire length of the array or absolute *DUXC* copy number in each cattle breed. Determining the length of the array is possible if there are ultra-long reads covering whole repeat units. Moreover, the individuals selected for the genomes may not represent the typical copy number of the *DUXC* unit.

## Conclusions

Our study establishes a comprehensive and experimentally validated annotation of bovine *DUXC*, revealing a novel upstream exon, confirming its functional domains, and elucidating its expression dynamics during early development. By uncovering the tandem repeat organization of the *DUXC* locus and demonstrating transcriptional activity of internal copies, we provide new insights into the structural complexity and evolutionary conservation of this gene across cattle breeds. These findings refine current models of embryonic genome activation and highlight *DUXC* as a candidate regulator of early developmental processes. Beyond improving genome annotation, our work lays the foundation for precise functional studies that will elucidate how repetitive homeobox loci contribute to species-specific regulation of zygotic genome activation.

## Methods

### PCR primer design

Putative protein sequence of porcine DUXC [27] was used to perform tblastn search [59] against the refseq_reference_genomes database from *Bos taurus* (taxid:9913) to find the *DUXC* locus. In this genomic locus, ENSBTAT00000081668, ENSBTAT00000070667, and ENSBTAT00000080304 from Ensembl release 104 and ENSBTAT00000111847 from Ensembl release 112 were selected as template sequences for PCR. Template sequences (DUXC_I-III) were extended toward the 5’ genomic region beyond the putative start codon to capture the 5’ UTRs. The template sequences (DUXC_IV f1/r1-f4/r4) for ENSBTAT00000111847 were extended toward the TFEs in the genomic region to allow the identification of potential novel upstream exons. The template sequences were extended also in the 3’ end to allow capturing the sequence beyond the putative stop codon. All template sequences were used as inputs for NCBI Primer-BLAST tool [60], in which *Bos taurus* (taxid:9913) was selected for primer pair specificity and repeat region was avoided. Forward and reverse primers with the lowest possible self-complementarity were chosen (Supplementary Table 2).

### Embryo culture, cDNA preparation and molecular cloning of *DUXC*

The 8-cell bovine IVF embryos were cultured and collected as described earlier [32]. cDNA library was prepared using the same protocol as in our previous study [32]. Similarly, for the cloning of *DUXC*, the same protocol described earlier [32] was used with the exception of minor modifications to PCR. Briefly, the PCR reaction was performed using Q5® High-Fidelity DNA Polymerase (New England Biolabs) following the protocol from the manufacturer using an input of 1 µl from the prepared cDNA library. PCR reactions were performed with betaine at a final concentration of 1M or without betaine. The parameters for PCR cycles were as follows: an initial denaturation at 98°C for 3 min, followed by 35 cycles of denaturation at 98°C for 10 s, annealing at 66°C (with betaine) or 68°C (without betaine) for 20 s, and extension at 72°C for 105 s, with a final extension at 72°C for 5 min. Three separate cloning experiments were conducted, where the same batch of pooled embryos were used for two of them, whereas a different batch was used for the other.

### Genomic organization analysis of *DUXC* loci in Hereford and Holstein bovine breeds

Chromosome 7 from Hereford (ARS-UCD2.0) and Holstein (NCBA_BosT1.0) [33] genome assemblies were extracted as FASTA files, which were converted to 2bit format using the UCSC faToTwobit utility [34]. Standalone in silico PCR program (isPcr, v33x2) [34] was utilized using the 2bit files along with forward (CCCGTGCCCCAGACCCC) and reverse (CCGGGCCCGTGGAAGCCCACTGGCT) primers with the options -out=fa -maxSize=7000 to perform in silico PCR. The primer pair was designed to anneal in opposite directions immediately adjacent to each other, ensuring that the entire length of each repeat unit was captured. Output FASTA files of in silico PCR were aligned separately for each breed using MAFFT (v.7.505) [61] with the flags --localpair --maxiterate 1000. Phylogenetic trees were built using IQ-TREE 2 (v.2.1.3) [62]. The packages ape (v.5.8.1) [63] and ggtree (v.3.14.0) [64] in R (v.4.4.1) were utilized for visualizing the resulting tree files.

### Extraction of *DUXC* loci and its annotation in multiple cattle breeds

Long-read sequencing-based genome assemblies of nine cattle breeds (Jersey [41], Hanwoo [41], Holstein [33], Hereford, Tibetan, Mongolian, Tuli, Wagyu [43] and Yunling) were downloaded from the NCBI, except for Holstein, which was downloaded from the China National Center for Bioinformation (Supplementary Table 6). Chromosome 7 from each breed was extracted using samtools (v.1.19) [65]. Internal *DUXC* repeat unit was used to perform alignments against the entire chromosome 7 from all breeds using minimap2 (v.2.26) [66] with the options -x asm20 -c --eqx after generating minimap2 indexes for chromosome 7 from each breed using the option -d. Outputs from the alignments were used to extract the entire array, with a 6 kb flanking sequencing added upstream and downstream of the array. First, each extracted *DUXC* locus from a given breed was used to perform alignments against the extracted *DUXC* locus of the other breeds, resulting in 36 pairwise alignment using minimap2 with the options -x asm20 -c --eqx. The resulting Pairwise mApping Format (paf) files were merged to be visualized by SVbyEye (v.0.99.0) [44]. Second, the internal *DUXC* repeat unit (Unit 2 from Holstein) was aligned against the extracted *DUXC* locus of each breed using minimap2 (-x asm20 -c --eqx). After creating minimap2 indexes for each extracted locus with the -d option, the full-length *DUXC* cDNA sequence was aligned to each locus using splice-aware alignment with -ax splice -N 40 options to create gene annotations. The resulting SAM files were converted to BAM and sorted using samtools (v.1.19), and then processed by StringTie (v.2.2.1) [38] to produce GTF files. The script agat_convert_sp_gxf2gxf.pl from AGAT [67] was used to convert GTF files to GFF3 format, which were used for visualization in R (v.4.4.1) by SVbyEye [44], together with the repeat unit versus *DUXC* locus alignments.

### Visualization of gene models, their primers and the structure of *DUXC* cDNA

Integrative Genomics Viewer (v.2.9.2) [68], UCSC Genome Browser [34] and R packages Gviz (v.1.48.0) [69], rtracklayer (v.1.64.0) [70], and GenomicRanges (v.56.1) [71] were utilized to visualize genomic regions. For visualization of Ensembl transcripts from release 104, GTF files for ENSBTAT00000081668, ENSBTAT00000070667, ENSBTAT00000080304 were retrieved through UCSC Table Browser [72], which were converted to GFF3 files using agat_convert_sp_gxf2gxf.pl script of the AGAT toolkit (v.1.4.2) [67]. For visualization of the Ensembl gene ENSBTAG00000049205 from release 112 using its corresponding transcripts ENSBTAT00000111847, ENSBTAT00000070667, and ENSBTAT00000081668, the Genepred format annotation file generated in our recent publication [26] by the STRTN_bTau_ens112.sh script (https://github.com/baryasar/bTau_Embryo_TFE) was utilized. For visualization of transcripts ENSBTAT00000081668, ENSBTAT00000070667, ENSBTAT00000080304 together with their primers, custom-generated primer BED files and GFF3 files were used. For visualizing ENSBTAT00000111847 and its primers, FASTA file of this transcript was used to map against the ARS-UCD1.2 genome assembly using the UCSC Blat tool [73], creating genomic coordinates compatible with accompanying BED file for the primers. Following mapping, UCSC Table Browser and AGAT (v.1.4.2) [67] were used to extract its GTF file and convert it to GFF3, respectively. BED file containing only *DUXC* TFEs were extracted from the TFE peak file generated earlier [26] and was lifted over to the ARS-UCD1.2 genome assembly. For visualizing the *DUXC* cDNA and its homeobox, a genome index was first created using gmap_build with the bovine genome assembly ARS-UCD1.3, and the corresponding FASTA files were then mapped with gmap (version 2018-07-04) to generate GFF outputs [39]. *DUXC* cDNA GFF3 file was converted to GenePred format using gff3ToGenePred utility [34]. For marking the 9aaTAD, a custom BED file was created. For visualizing *DUXC* cDNA at the extracted internal and distal regions gmap_build was used to index each region separately, to which *DUXC* cDNA FASTA file was aligned using gmap (version 2018-07-04) to generate GFF3 annotation files [39].

### Pairwise and multiple sequence alignment

EMBOSS Water tool via EMBL-EBI Job Dispatcher [74] was used to perform pairwise alignment of the putative exon 5 from the distal and internal units. For multiple sequence alignment, exon 1 FASTA sequence was obtained from breeds with annotated *DUXC* by filtering the GFF3 annotation files for the gene_id DUXC.1 using the agat_sp_filter_feature_from_keep_list.pl script from AGAT (v.1.4.2) and then extracting the sequences by Gffread (v.0.12.8) [75]. Clustal Omega via EMBL-EBI Job Dispatcher was utilized to perform multiple sequence alignment of exon 1 extracted from Hanwoo, Hereford, Holstein, Tibetan, Tuli, Wagyu and Jersey cattle breeds.

### Identification of telomeric repeats in Holstein chromosome 7

Holstein genome assembly [33] was used as an input for tidk explore function (v.0.2.65) with the flags --minimum 5 --maximum 12 --distance 0.01 [40]. Five most frequent motifs were used to execute tidk explore and tidk plot function on chromosome 7.

### Identification of polyadenylation signals

A search in the 3’-UTRs of *DUXC* and its downstream regions in Holstein (ARS-UCD2.0) and Hereford (NCBA_BosT1.0) cattle was performed using the 12 main polyadenylation signal variants [76].

### Reanalysis of RNA-seq datasets

For assessing the expression of putative distal *DUXC*, paired-end RNA-seq reads from a published study [36] were utilized. Distal and internal units from Holstein cattle chromosome 7 (NCBA_BosT1.0) were extracted, with their boundaries extended upstream beyond the repeat units borders defined in this study to ensure that exon 1 was included, which spans the end of one unit and the beginning of the other. Using HISAT2 (v.2.2.1) [19] with default parameters, RNA-seq reads were aligned to these genomic regions that contain full-length sequences of putative distal and internal *DUXC*. The resulting SAM files were converted to BAM format using samtools (v.1.6) [65] with the option -F 4 to retain only mapped reads followed by sorting and indexing. Alignment files were merged based on the stage using samtools merge and indexed. StringTie (v.2.1.7) [38] was used for transcript assembly for each stage. For annotating homeodomain-encoding sequences, gmap (version 2018-07-04) was used to map homeobox sequences in the internal and distal *DUXC* to internal and distal *DUXC* genomic locus, respectively. Integrative Genomics Viewer (v.2.9.2) [68] and R package Gviz (v.1.48.0) [69] were used to visualize the alignment files.

For the reanalysis of the RNA-seq dataset from a recent study [31], ARS-UCD1.3 genome assembly and Ensembl gene annotation file ARS-UCD1.3.112 were used. Integrating the cDNA sequences of *DUXC* and seven other PRDL genes [32] in the bovine annotation was performed as previously described [26]. A portion of chromosome 7 was masked using bedtools maskfasta (v.2.30.0) [77] to ensure mapping of RNA-seq reads to only one copy of *DUXC*, which left the genomic region between the nucleotides 85,000 and 92,000 unmasked. Splice sites and exons were extracted and used for building the genome index with subsequent mapping using HISAT2 (v.2.2.1) [19]. The resulting SAM files converted to BAM files and sorted and indexed using samtools (v.1.6) [65]. Quantification of expression was performed at the gene-level using featureCounts (v.2.0.1) [78]. The resulting gene count table was imported into R (v.4.4.1) and a DESeqDataSet using DESeq2 (v.1.44.0) [79] was formed by setting the developmental stage in the design formula. Genes with less than 10 reads summed across all samples were removed from DESeqDataSet. Next, variance-stabilizing transformation (VST) was performed, followed by z-score normalization for visualization using R packages pheatmap (v.1.0.13), RColorBrewer (v.1.1.3) and viridis (v.0.6.5) [80]. For the reanalysis of our recently published dataset, the same preprocessing and analysis pipeline was used, except for the additional masking step mentioned above [26]. Within this workflow, the STRTN_bTau_ens112.sh script compatible with the Ensembl gene annotation file was executed to generate a gene-based read count file. All computational work, except for the R scripts, was performed at the High Performance Computing Center at the University of Tartu [81].

## Declarations

### Ethics and approval and consent to participate

Not applicable.

### Consent for publications

Not applicable.

### Availability of data and material

The dataset generated during the current study is available in the European Nucleotide Archive (ENA) repository, https://www.ebi.ac.uk/ena/data/view/PRJEB104384.

### Authors’ contributions

B.Y.: Investigation, Validation, Data curation, Software, Formal analysis, Visualization, Writing – original draft; B.Y., S.K.: Conceptualization, Project administration; S.K., J.K., A.K. and T.O.: Resources, Supervision, Funding Acquisition; B.Y., T.O., M.I., G.Y.G., N.B., Ü.J., J.K., A.K., S.K.: Methodology, Writing – review & editing.

## Acknowledgements

The authors wish to thank Merle Külaots for her assistance with the plasmid extraction. The icon in Figure 3C was created with BioRender.com.

## Additional Files

### Additional file 1

**Supplementary Figure 1. (A)** UCSC Genome Browser (http://genome.ucsc.edu) screenshot of chromosome 7 from the ARS-UCD1.2 genome assembly, showing predicted *DUXC* gene structures (Ensembl Gene Predictions version 104). Genes with identical putative spliced transcript sequences are marked with stars of the same colour. **(B)** Uncropped image of the agarose gel in Fig. 1D. (.pdf)

**Supplementary Figure 2**. Integrative Genomics Viewer (IGV) screenshot of mapped RNA-seq reads derived from the 1-, 2-, 4-, 8-, and 16-cell stage embryos cultured under normal conditions (Zhang et al., 2025) to the ARS-UCD1.3 bovine genome assembly. (.pdf)

**Supplementary Figure 3**. Visualisation of polyadenylation signals detected in the downstream region of the 3’-UTRs of *DUXC* copies. (.pdf)

**Supplementary Figure 4**. Pairwise alignment of exon 5 from Holstein internal and distal repeat units. The putative distal *DUXC* exon 5 is extended towards the 3’ end to include genomic sequences in this region, allowing comparison with the internal counterpart. Red dashed vertical line indicates the region after which the sequence (to the left; towards 3’ end) starts to differ between putative distal and internal *DUXC*. (.pdf)

**Supplementary Figure 5**. Occurrence of the repeat “ATCATGC” and “ATCATGCATG”, which were explored by tidk in chromosome 7 of Holstein genome assembly. The peak at the left end of the chromosome indicates the assembled telomere at this end. (.pdf)

**Supplementary Figure 6**. Integrative Genomics Viewer (IGV) screenshot of the alignments of the 2-cell stage RNA-seq reads (Zhu et al., 2022) to the **(A)** distal and **(B)** internal genomic region of Holstein chromosome 7. (.pdf)

**Supplementary Figure 7**. Alignment of RNA-seq reads derived from the **(A)** GV oocyte, **(B)** MII oocyte, **(C)** morula and **(D)** blastocyst stage using a publicly available dataset (Zhu et al., 2022) to the distal (Unit 1) and internal (Unit 2) *DUXC* units of Holstein cattle. (.pdf)

**Supplementary Figure 8**. Holstein internal *DUXC* repeat unit (Unit 2) alignment against the entire *DUXC* locus of **(A)** Jersey, **(B)** Hanwoo, **(C)** Mongolian, **(D)** Tuli, **(E)** Tibetan, and **(F)** Wagyu cattle. (.pdf)

**Supplementary Figure 9**. Clustal Omega multiple sequence alignment of *DUXC* exon 1 across cattle breeds. (.pdf)

### Additional file 2

**Supplementary Table 1**. Known binders of the de novo motifs identified in the promoters of upregulated 16-cell-specific transcripts. (.xlsx)

**Supplementary Table 2**. PCR primers used for the amplification of *DUXC* cDNA. (.xlsx)

**Supplementary Table 3**. GC content of expected PCR products. (.xlsx)

**Supplementary Table 4**. In silico PCR results. (.xlsx)

**Supplementary Table 5**. Genomic coordinates of the extracted *DUXC* locus in eight cattle breeds. (.xlsx)

**Supplementary Table 6**. Long-read sequencing-based genome assemblies and their associated information, including contig N50 lengths and genome coverage, used in the comparative analysis of the *DUXC* locus. (.xlsx)

**Supplementary Table 7**. Genomic coordinates of each *DUXC* repeat unit in eight cattle breeds. (.xlsx)

### Additional file 3

**Supplementary Text 1**. Distal *DUXC* repeat units of Hereford and Holstein called Unit 1. (.pdf)

**Supplementary Text 2**. Final *DUXC* repeat units of Hereford and Holstein called Unit 8 and Unit 20, respectively. (.pdf)

**Supplementary Text 3**. Unit 2 that was used to extract *DUXC* loci from each breed. (.pdf)

