## Supplementary_figures for "Structure and genomic organization of the human *DUX4* homologue bovine *DUXC*"

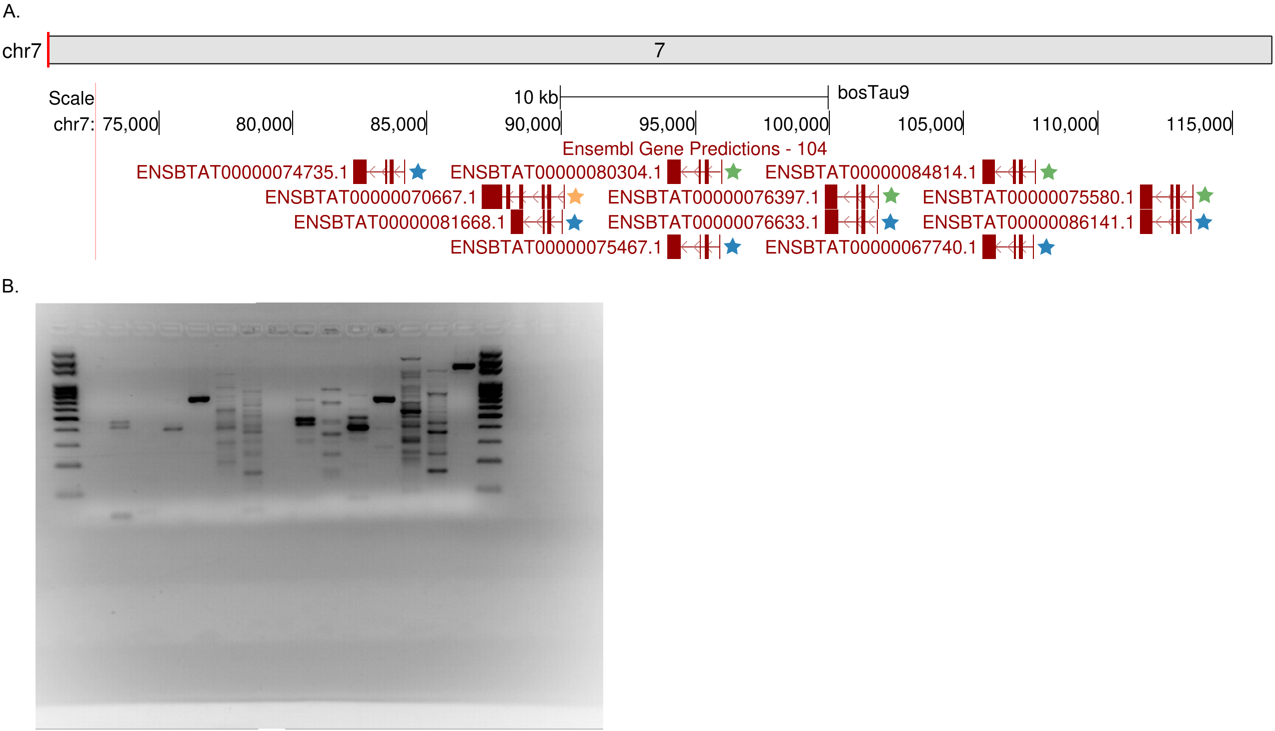


**Supplementary Figure 1. (A)** UCSC Genome Browser (http://genome.ucsc.edu) screenshot of chromosome 7 from the ARS-UCD1.2 genome assembly, showing predicted *DUXC* gene structures (Ensembl Gene Predictions version 104). Genes with identical putative spliced transcript sequences are marked with stars of the same color. **(B)** Uncropped image of the agarose gel in Fig. 1D.

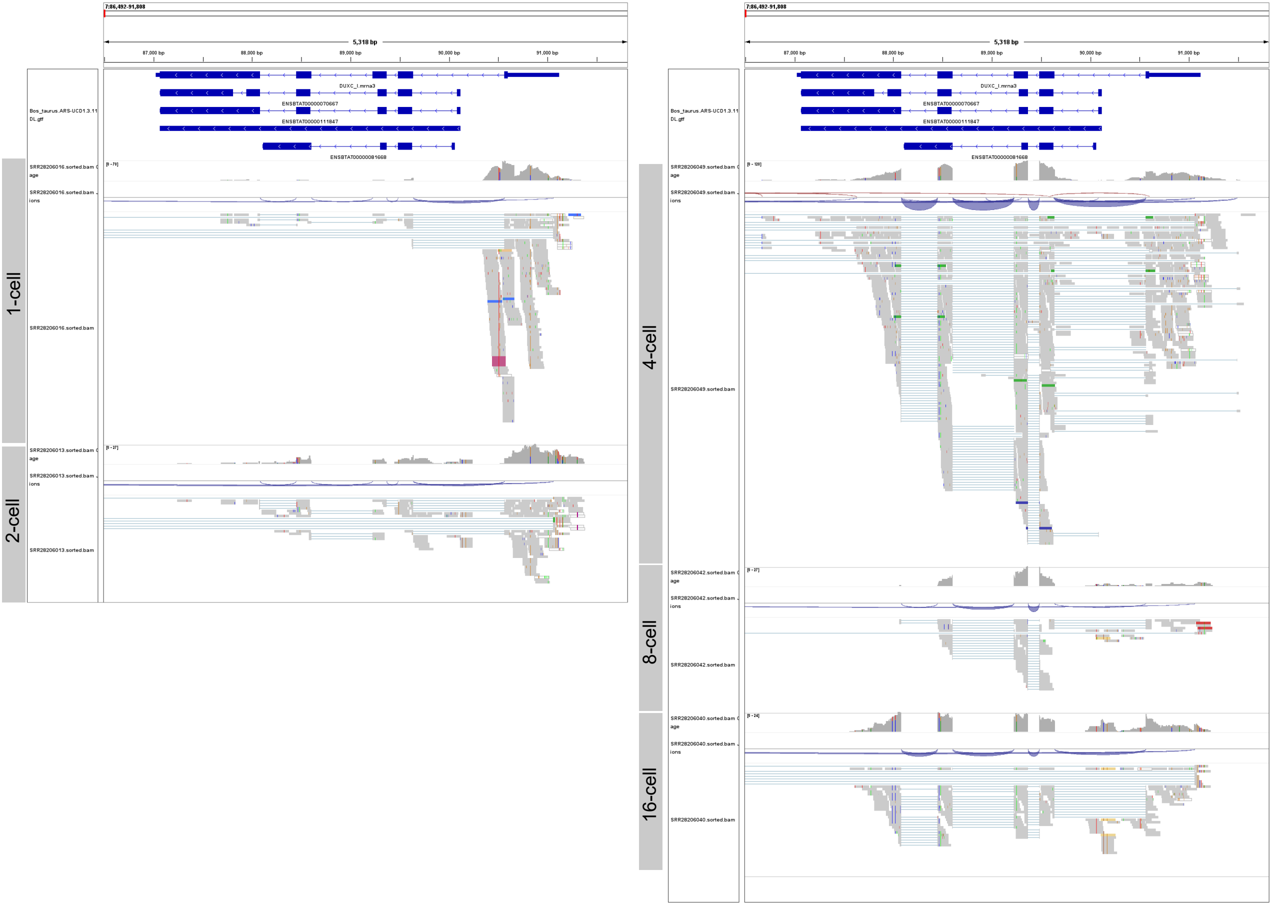


**Supplementary Figure 2.** Integrative Genomics Viewer (IGV) screenshot of mapped RNA-seq reads derived from the 1-, 2-, 4-, 8-, and 16-cell stage embryos cultured under normal conditions (Zhang et al., 2025) to the ARS-UCD1.3 bovine genome assembly.

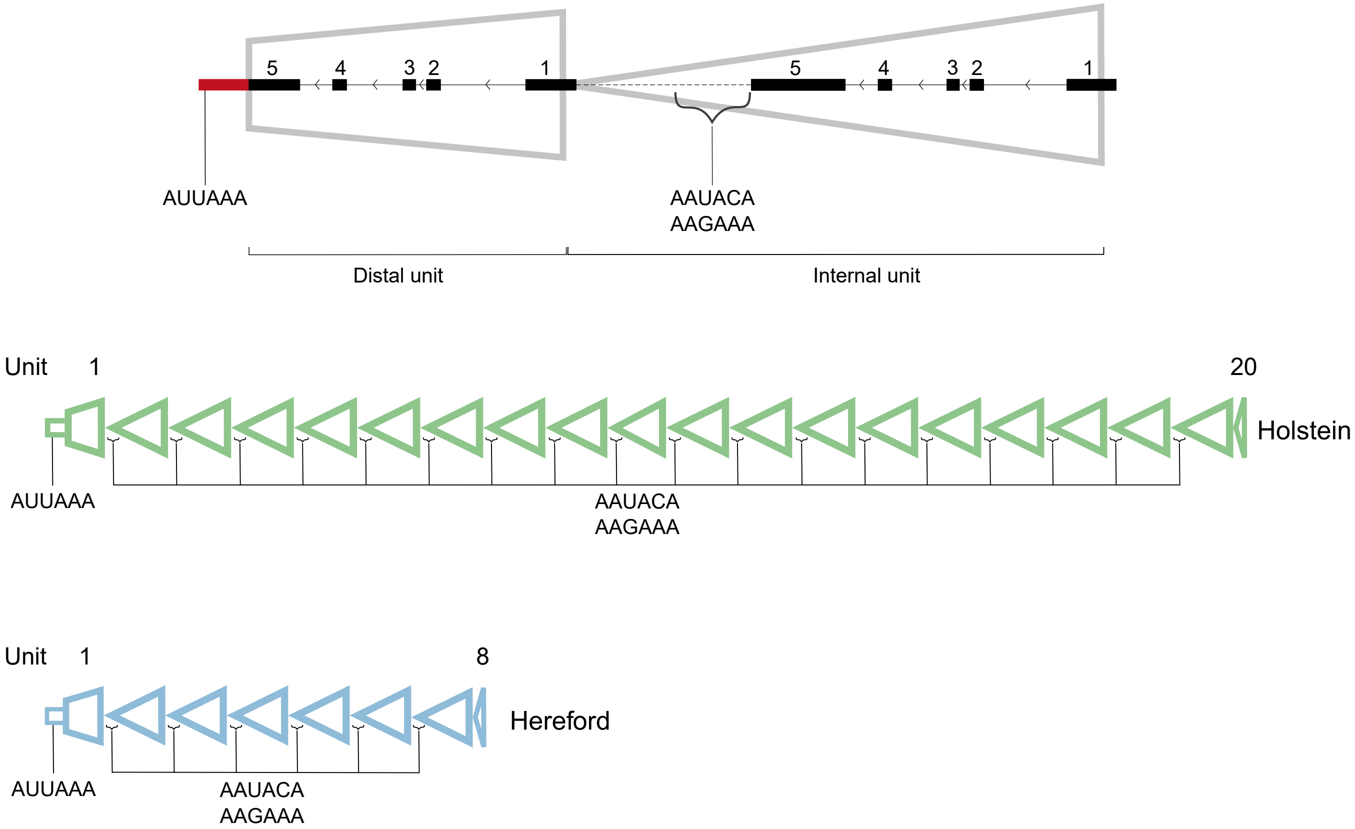


**Supplementary Figure 3.** Visualisation of polyadenylation signals detected in the downstream region of the 3′-UTRs of *DUXC* copies.

```
distalExon5        8 GGCAAGGGTAG----GAA-GTGG-TAGTGGGATGATTGGCTGGGTTTCCC     51
                     |||||||  ||    ||| |||| |||.||||..|       |||..|||
internalExon5    101 GGCAAGG--AGACCTGAACGTGGCTAGGGGGAGAA-------GGTGGCCC    141

distalExon5       52 ACACTTCTCAGTGCTT--TAATGAAACACCCTCCTGGCTTAATCACC---     96
                        ||||      |||  |       |.|||||      |||.|.||
internalExon5    142 ---CTTC------CTTCCT-------CTCCCTC------TAAGCCCCAGG    169

distalExon5       97 TCCTGGGCATTCTGGTTCA-CATG----------------------CCTC    123
                     .||||||.|..||||  || ||||                      ||||
internalExon5    170 GCCTGGGGAGCCTGG--CAGCATGTCGAGGAGGGCCTGGTACTCGTCCTC    217

distalExon5      124 AAT-----GTGTCACATCCTGGGACTAT-GTCTCAGCT--ATCTT-----    160
                     |.|     ||||   .|||.|||    | |.||||.||  .||||
internalExon5    218 ACTGAGGGGTGT---TTCCAGGG----TGGCCTCAACTCCTTCTTCCCCC    260

distalExon5      161 ----GGGACTATCCCCCAGCTCACATCATGGCATGCCTGTTCCCATGTGT    206
                         ||||     ||||.|...|.|.||.|||              ||||
internalExon5    261 GCAGGGGA-----CCCCGGGGAAGACCAGGGC--------------GTGT    291

distalExon5      207 TCACTCCGCTACCTGGTTGATCCTGTCAG-TAATGTTTGAAACATTAAGC    255
                                |||||.||..|.||.||| .||..|||
internalExon5    292 -----------CCTGGATGGCCGTGGCAGCCAAGATTT------------    318

distalExon5      256 AGAGCATGTCTAAAGAACACCT---GGTACAGGAAAACTTGAAAAGGGTG    302
                                        |||   ||   |||    ||      |||||
internalExon5    319 -------------------CCTCTAGG---AGG----CT------GGGTG    336

distalExon5      303 TG--------GTGCTGGGCTCTGCTGGGG---TGCTGAACTCTACAGTTT    341
                      |        |.||||||      |||||   ||| ||   |.||||
internalExon5    337 -GCCCGGGGAGGGCTGGG------TGGGGCCCTGC-GA---CGACAG---    372

distalExon5      342 TGAAACCATATCTCTAGG-----GGAGCAAAGGTAGGTAGGCACATCAGA    386
                      ||           .|||     ||||  |||       |||.|..| ||
internalExon5    373 -GA-----------GAGGACCCCGGAG--AAG-------GGCCCGGC-GA    400

distalExon5      387 ACATGTGATGTCC------------AATTCACC--------------ATG    410
                     .|..|.||.|.||            |..||.||              .||
internalExon5    401 GCCCGGGAGGCCCGCGGCAGCCAGGAGCTCGCCCGGGAGGGCGAGGTGTG    450

distalExon5      411 CTGGTGCCTCTGAAGCAACTATAAGAAAGCCTGCATGGAAGATTGGCCTG    460
                     ||.||.|.||.|.||           |.|.|..||.|||||   |.||.|
internalExon5    451 CTCGTTCATCGGCAG-----------AGGGCCCCAGGGAAG---GCCCCG    486

distalExon5      461 TGGTCTGTAAGGCCCAGGGTGATGGTAGGTAGGCAG-CATGAG-----CC    504
                     ..||       |.||.||||||...||    |||.| ||.|||     |.
internalExon5    487 GCGT-------GTCCGGGGTGCCCGT-----GGCCGCCAAGAGCTCGTCT    524

distalExon5      505 AGGAA-CT-----CCCCTGGGAGGATG-AGGTGTG----CTCGTTCATTG    543
                     ||||| ||     |||||||||||  | |||   |    |.|.|||.|.|
internalExon5    525 AGGAAGCTGGGTGCCCCTGGGAGG--GCAGG---GCCCACCCCTTCGTCG    569

distalExon5      544 GCAGCGGCCCCCAGGGAAGGCGCCTGCGTGTCCGGGACGCCCGTGGCCGC    593
                     ||||||||||||.|||||||||||||||||||||||||||||||||||||
internalExon5    570 GCAGCGGCCCCCGGGGAAGGCGCCTGCGTGTCCGGGACGCCCGTGGCCGC    619

distalExon5      594 CAAGAGCTCATCTAGGAGGCTCGTCGAGCTTGGAAGGCTGGGGGCCTGGG    643
                     ||||||||||||||||||||||||||||||||||||||||||||||||||
internalExon5    620 CAAGAGCTCATCTAGGAGGCTCGTCGAGCTTGGAAGGCTGGGGGCCTGGG    669

distalExon5      644 GCTGCGGCTCCAGCGGAGGGGCCGTGCCTTCGCCGGAGGGTGACTCCGGC    693
                     ||||||||||||||||||||||||||||||||||||||||||||||||||
internalExon5    670 GCTGCGGCTCCAGCGGAGGGGCCGTGCCTTCGCCGGAGGGTGACTCCGGC    719

distalExon5      694 CACCGCGGGATGTGTGTCTCCGGCTGCCCAGGCGGCAGGATGGCCCCTTG    743
                     ||||||||||||||||||||||||||||||||||||||||||||||||||
internalExon5    720 CACCGCGGGATGTGTGTCTCCGGCTGCCCAGGCGGCAGGATGGCCCCTTG    769

distalExon5      744 CCCAGGCAGCCCGGGGGCCGTGACGGCAGGGGAGCACGCGGCCCAGGGAG    793
                     ||||||||||||||||||||||||||||||||||||||||||||||||||
internalExon5    770 CCCAGGCAGCCCGGGGGCCGTGACGGCAGGGGAGCACGCGGCCCAGGGAG    819

distalExon5      794 CTGCTCCCTGGGGAGGAGGCGGTGGCCTCCCACTTGGCTGGAGGGCCGCC    843
                     ||||||||||||||||||||||||||||||||||||||||||||||||||
internalExon5    820 CTGCTCCCTGGGGAGGAGGCGGTGGCCTCCCACTTGGCTGGAGGGCCGCC    869

distalExon5      844 GGGCTGGGCTGGACCACGAAGATCATCAACGGCTGGCCCACGCAGACCCC    893
                     ||||||||||||||||||||||||||||||||||||||||||||||||||
internalExon5    870 GGGCTGGGCTGGACCACGAAGATCATCAACGGCTGGCCCACGCAGACCCC    919

distalExon5      894 AGAGGCAGCCCCGGGCACCCAGAAAGTGGGGGCCCCAAAAGGAGCAGCTG    943
                     ||||||||||||||||||||||||||||||||||||||||||||||||||
internalExon5    920 AGAGGCAGCCCCGGGCACCCAGAAAGTGGGGGCCCCAAAAGGAGCAGCTG    969

distalExon5      944 CAGCCAAGGGTGGCATGCTCTCCTGTGGCTGAGAGGGGCGAAGGGGAGGA    993
                     ||||||||||||||||||||||||||||||||||||||||||||||||||
internalExon5    970 CAGCCAAGGGTGGCATGCTCTCCTGTGGCTGAGAGGGGCGAAGGGGAGGA   1019

distalExon5      994 GAGGTGCTCTGGACAGCGGGTGGGGCCCTTCGGTCCTCTGGAGCAGGAGT   1043
                     ||||||||||||||||||||||||||||||||||||||||||||||||||
internalExon5   1020 GAGGTGCTCTGGACAGCGGGTGGGGCCCTTCGGTCCTCTGGAGCAGGAGT   1069

distalExon5     1044 CGTCGTCGTGGCTGGTGCGCCGCCTGGCCCCTGGGCCCGGCCGTTGCCGG   1093
                     ||||||||||||||||||||||||||||||||||||||||||||||||||
internalExon5   1070 CGTCGTCGTGGCTGGTGCGCCGCCTGGCCCCTGGGCCCGGCCGTTGCCGG   1119

distalExon5     1094 GCCCGCTTGGG   1104
                     |||||||||||
internalExon5   1120 GCCCGCTTGGG   1130
```

**Supplementary Figure 4.** Pairwise alignment of exon 5 from Holstein internal and distal repeat units. The putative distal *DUXC* exon 5 is extended towards the 3' end to include genomic sequences in this region, allowing comparison with the internal counterpart. The red dashed vertical line indicates the region after which the sequence (to the left; towards 3' end) starts to differ between putative distal and internal *DUXC*.

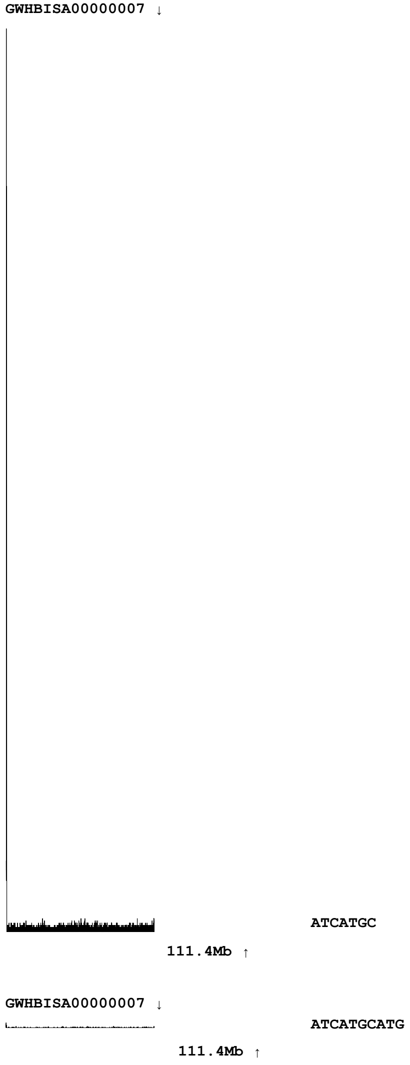


**Supplementary Figure 5.** Occurence of the repeat "ATCATGC" and "ATCATGCATG", which were explored by tidk in chromosome 7 of Holstein genome assembly. The peak at the left end of the chromosome indicates the assembled telomere at this end.

A.

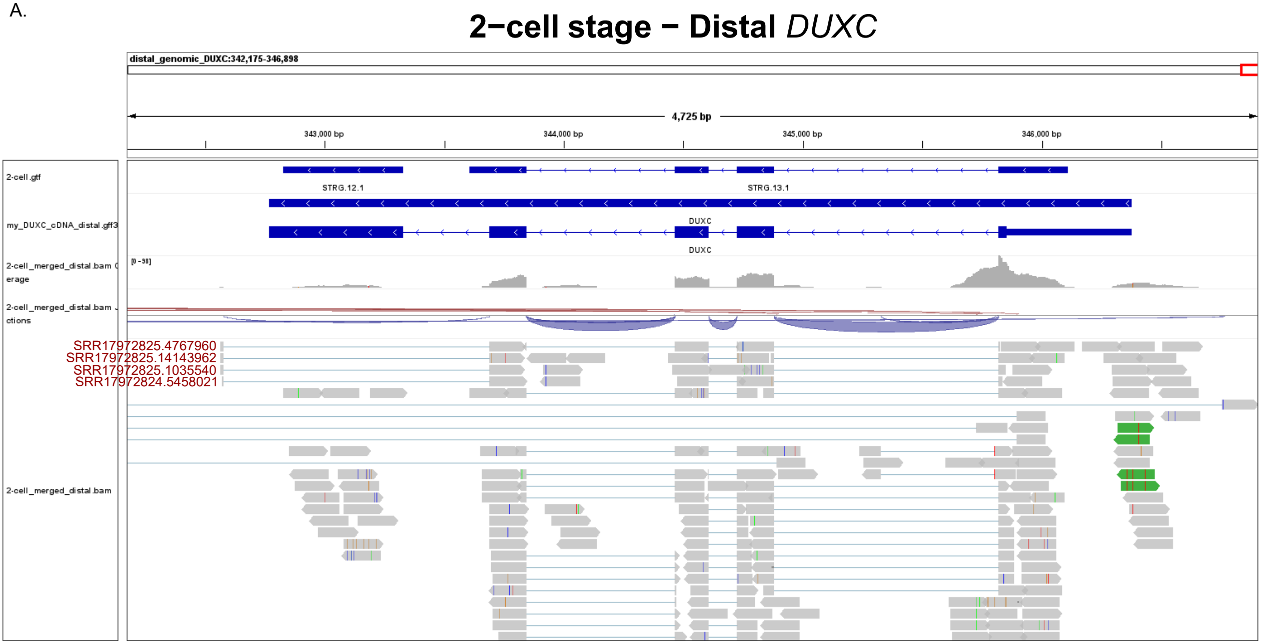


B.

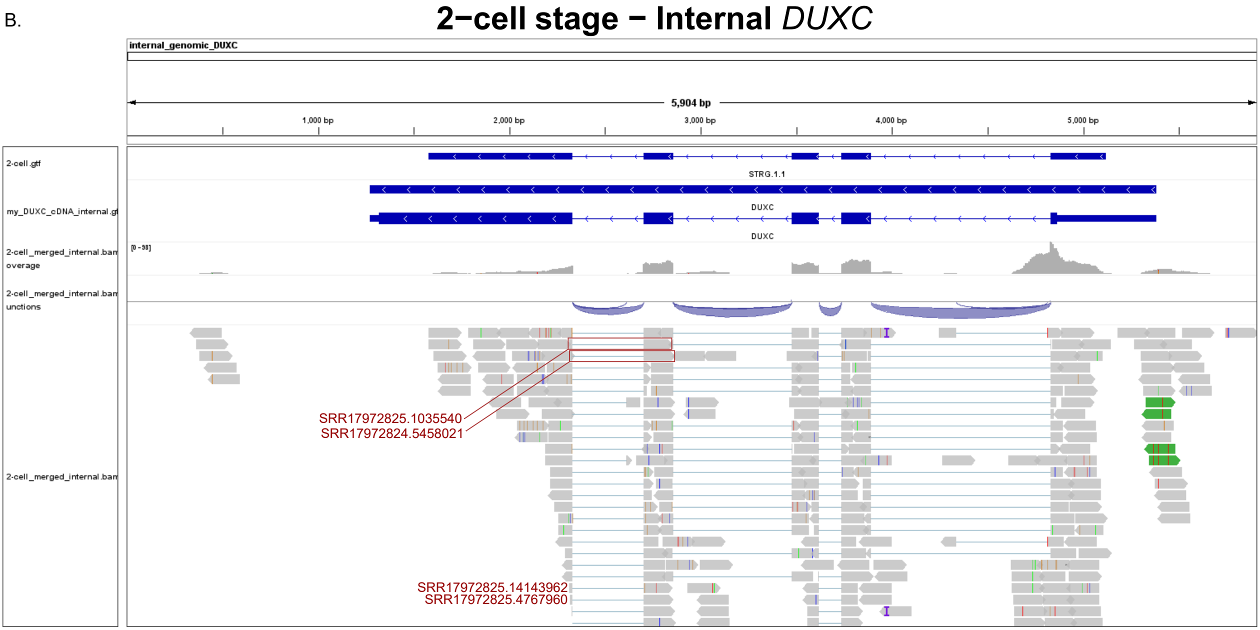


**Supplementary Figure 6.** Integrative Genomics Viewer (IGV) screenshot of the alignments of the 2-cell stage RNA-seq reads (Zhu et al., 2022) to the **(A)** distal and **(B)** internal genomic region of Holstein chromosome 7.

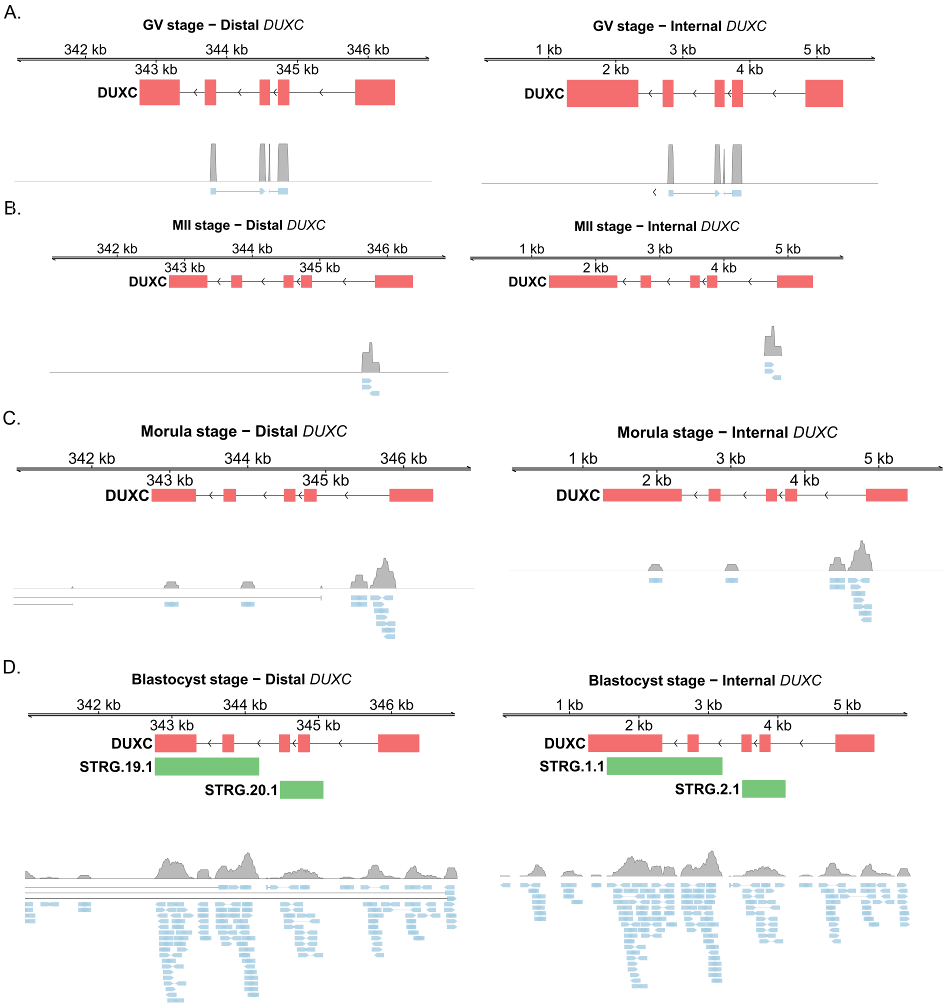


**Supplementary Figure 7.** Alignment of RNA-seq reads derived from the **(A)** GV oocyte, **(B)** MII oocyte, **(C)** morula and **(D)** blastocyst stage using a publicly available dataset (Zhu et al., 2022) to the distal (Unit 1) and internal (Unit 2) *DUXC* units of Holstein cattle.

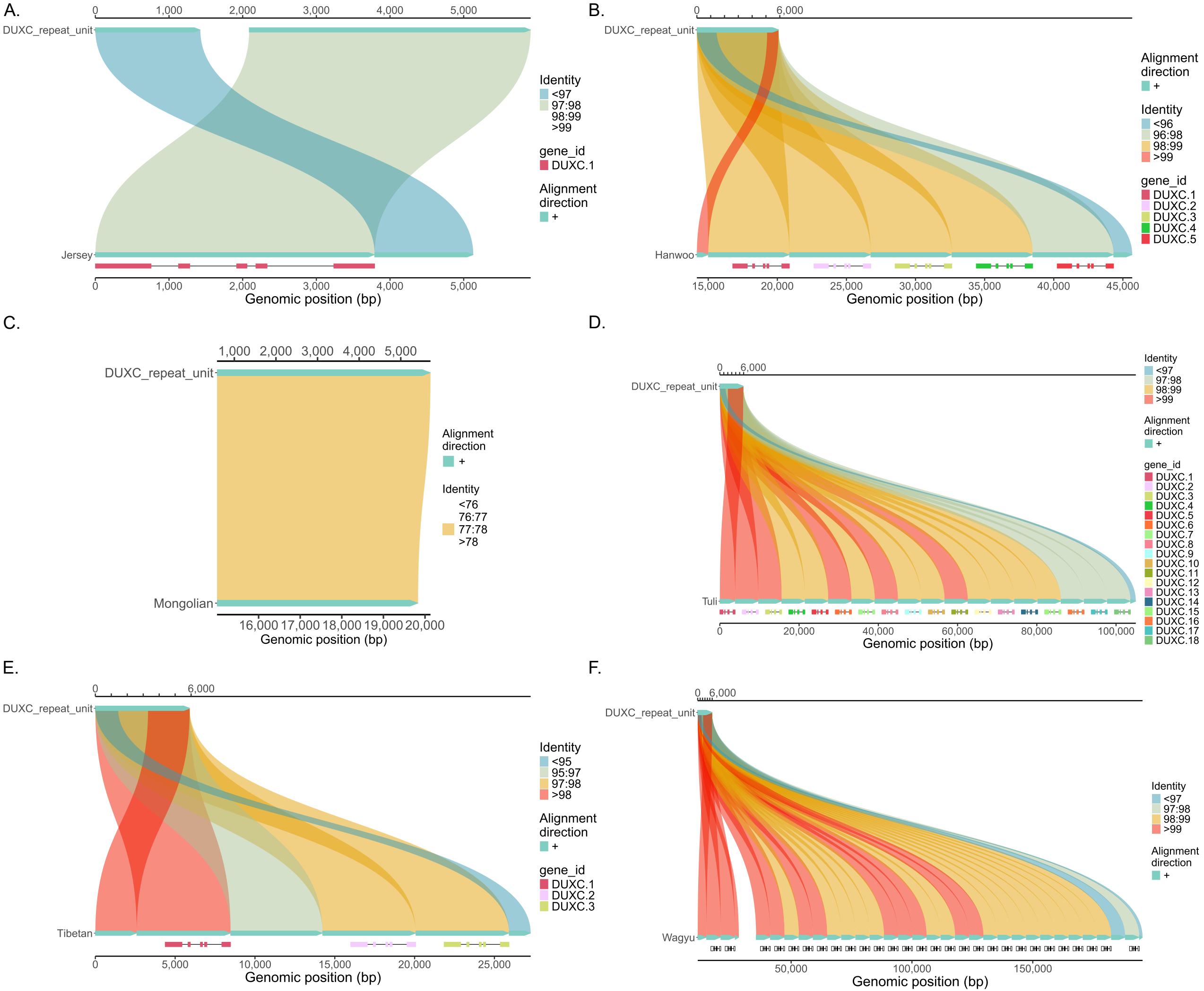


**Supplementary Figure 8.** Holstein internal *DUXC* repeat unit (Unit 2) alignment against the entire *DUXC* locus of **(A)** Jersey, **(B)** Hanwoo, **(C)** Mongolian, **(D)** Tuli, **(E)** Tibetan, and **(F)** Wagyu cattle.

```
Hanwoo_DUXC.1.1        GCATTGGAGAGAGCATCGGGTGTCCCGCCATGCCTCGCAGGAGCCGGCGGGGGCAATTGA 60
Hereford_DUXC.1.1      GCATTGGAGAGAGCATCGGGTGTCCCGCCATGCCTCGCAGGAGCCGGCGGGGGCAATTGA 60
Holstein_DUXC.1.1      GCATTGGAGAGAGCATCGGGTGTCCCGCCATGCCTCGCAGGAGCCGGCGGGGGCAATTGA 60
Tibetan_DUXC.1.1       GCATTGGAGAGAGCATCGGGTGTCCCGCCATGCCTCGCAGGAGCCGGCGGGGGCAATTGA 60
Tuli_DUXC.1.1          GCATTGGAGAGAGCATCGGGTGTCCCGCCATGCCTCGCAGGAGCCGGCGGGGGCAATTGA 60
Wagyu_DUXC.1.1         GCATTGGAGAGAGCATCGGGTGTCCCGCCATGCCTCGCAGGAGCCGGCGGGGGCAATTGA 60
Jersey_DUXC.1.1        GCATTGGAGAGAGCATCGGGTGTCCCGCCATGCCTCGCAGGAGCCGGCGGGGGCAATTGA 60
                       ************************************************************

Hanwoo_DUXC.1.1        GGTCAAGGGACGTGTGGAATGTGAAAAGGCGGGAAAACCCAGAGCGGCTTGGAAGGCAAG 120
Hereford_DUXC.1.1      GGTCAAGGGACGTGTGGAATGTGAAAAGGCGGGAAAACCCAGAGCGGCTTGGAAGGCAAG 120
Holstein_DUXC.1.1      GGTCAAGGGACGTGTGGAATGTGAAAAGGCGGGAAAACCCAGAGCGGCTTGGAAGGCAAG 120
Tibetan_DUXC.1.1       GGTCAAGGGACGTGTGGAATGTGAAAAGGCGGGAAAACCCAGAGCGGCTTGGAAGGCAAG 120
Tuli_DUXC.1.1          GGTCAAGGGACGTGTGGAATGTGAAAAGGCGGGAAAACCCAGAGCGGCTTGGAAGGCAAG 120
Wagyu_DUXC.1.1         GGTCAAGGGACGTGTGGAATGTGAAAAGGCGGGAAAACCCAGAGCGGCTTGGAAGGCAAG 120
Jersey_DUXC.1.1        GGTCAAGGGACGTGTGGAATGTGAAAAGGCGGGAAAACCCAGAGCGGTTTGGAAGGCAAG 120
                       *********************************************** ************

Hanwoo_DUXC.1.1        GCAGGCTGGGCAAGGGGGCGTGGCCGGCCACGTCAAAGGGACCCCAGGGGAATTCCCGGG 180
Hereford_DUXC.1.1      GCAGGCTGGGCAAGGGGGCGTGGCCGGCCACGTCAAAGGGACCCCAGGGGAATTCCCGGG 180
Holstein_DUXC.1.1      GCAGGCTGGGCAAGGGGGCGTGGCCGGCCACGTCAAAGGGACCCCAGGGGAATTCCCGGG 180
Tibetan_DUXC.1.1       GCAGGCTGGGCAAGGGGGCGTGGCCGGCCACGTCAAAGGGACCCCAGGGGAATTCCCGGG 180
Tuli_DUXC.1.1          GCAGGCTGGGCAAGGGGGCGTGGCCGGCCACGTCAAAGGGACCCCAGGGGAATTCCCGGG 180
Wagyu_DUXC.1.1         GCAGGCTGGGCAAGGGGGCGTGGCCGGCCACGTCAAAGGGACCCCAGGGGAATTCCCGGG 180
Jersey_DUXC.1.1        GCAGGCTGGGCAAGGGGGCGTGGCCGGCCACGTCAAAGGGACCCCAGGGGAATTCCCGGG 180
                       ************************************************************

Hanwoo_DUXC.1.1        CAGTTAGCTCTCTGGGCACTGCTGACGGCCCCTGGGCCCGGGGTTGACCCCACCCGCTGC 240
Hereford_DUXC.1.1      CAGTTAGCTCTCTGGGCACTGCTGACGGCCCCTGGGCCCGGGGTTGACCCCACCCGCTGC 240
Holstein_DUXC.1.1      CAGTTAGCTCTCTGGGCACTGCTGACGGCCCCTGGGCCCGGGGTTGACCCCACCCGCTGC 240
Tibetan_DUXC.1.1       CAGTTAGCTCTCTGGGCACTGCTGACGGCCCCTGGGCCCGGGGTTGACCCCACCCGCTGC 240
Tuli_DUXC.1.1          CAGTTAGCTCTCTGGGCACTGCTGACGGCCCCTGGGCCCGGGGTTGACCCCACCCGCTGC 240
Wagyu_DUXC.1.1         CAGTTAGCTCTCTGGGCACTGCTGACGGCCCCTGGGCCCGGGGTTGACCCCACCCGCTGC 240
Jersey_DUXC.1.1        CCGTTAGCTCTCTGGGCACTGCTGACGGCCCCTGGGCCCGGGGTTGACCCCACCCGCTGC 240
                       * **********************************************************

Hanwoo_DUXC.1.1        GTGGGAGGGCCACAAACCAAAGCCCCCGAACTGCCCGCCCAGCCCAACGGTGGACCTTGG 300
Hereford_DUXC.1.1      GTGGGAGGGCCACAAACCAAAGCCCCCGAACTGCCCGCCCAGCCCAACGGTGGACCTTGG 300
Holstein_DUXC.1.1      GTGGGAGGGCCACAAACCAAAGCCCCCGAACTGCCCGCCCAGCCCAACGGTGGACCTTGG 300
Tibetan_DUXC.1.1       GTGGGAGGGCCACAAACCAAAGCCCCCGAACTGCCCGCCCAGCCCAACGGTGGACCTTGG 300
Tuli_DUXC.1.1          GTGGGAGGGCCACAAACCAAAGCCCCCGAACTGCCCGCCCAGCCCAACGGTGGACCTTGG 300
Wagyu_DUXC.1.1         GTGGGAGGGCCACAAACCAAAGCCCCCGAACTGCCCGCCCAGCCCAACGGTGGACCTTGG 300
Jersey_DUXC.1.1        GTGGGAGGGCCACATACCAAAGCCCCCGAACTGCCCGCCCAGCCCAACGGTGGACCTTGG 300
                       ************** *********************************************

Hanwoo_DUXC.1.1        GGGTCTGGGGCACGGGCCGGGCCCGTGGAAGCCCACTGGCTCCACTGGATTCTTGTCCAG 360
Hereford_DUXC.1.1      GGGTCTGGGGCACGGGCCGGGCCCGTGGAAGCCCACTGGCTCCACTGGATTCTTGTCCAG 360
Holstein_DUXC.1.1      GGGTCTGGGGCACGGGCCGGGCCCGTGGAAGCCCACTGGCTCCACTGGATTCTTGTCCAG 360
Tibetan_DUXC.1.1       GGGTCTGGGGCACGGGCCGGGCCCGTGGAAGCCCACTGGCTCCACTGGATTCTTGTCCAG 360
Tuli_DUXC.1.1          GGGTCTGGGGCACGGGCCGGGCCCGTGGAAGCCCACTGGCTCCACTGGATTCTTGTCCAG 360
Wagyu_DUXC.1.1         GGGTCTGGGGCACGGGCCGGGCCCGTGGAAGCCCACTGGCTCCACTGGATTCTTGTCCAG 360
Jersey_DUXC.1.1        GGGTCTGGGGCACGGGCCGGGCCCGTGGAAGCCCACTGGCTCCACTGGATTCTTGTCCAG 360
                       ************************************************************

Hanwoo_DUXC.1.1        GGTCTCTCCTGCGAAAGAGGCCAGGGGCGTCCCCTCCATCTACGCACCCACCCCTATTTA 420
Hereford_DUXC.1.1      GGTCTCTCCTGCGAAAGAGGCCAGGGGCGTCCCCTCCATCTACGCACCCACCCCTATTTA 420
Holstein_DUXC.1.1      GGTCTCTCCTGCGAAAGAGGCCAGGGGCGTCCCCTCCATCTACGCACCCACCCCTATTTA 420
Tibetan_DUXC.1.1       GGTCTCTCCTGCGAAAGAGGCCAGGGGCGTCCCCTCCATCTACGCACCCACCCCTATTTA 420
Tuli_DUXC.1.1          GGTCTCTCCTGCGAAAGAGGCCAGGGGCGTCCCCTCCATCTACGCACCCACCCCTATTTA 420
Wagyu_DUXC.1.1         GGTCTCTCCTGCGAAAGAGGCCAGGGGCGTCCCCTCCATCTACGCACCCACCCCTATTTA 420
Jersey_DUXC.1.1        GGTCTCTCCTGCGAAAGAGGCGAGGGGCGTCCCCTCCATCTACGCACCCACTCCTATTTA 420
                       ********************* ***************************** ********

Hanwoo_DUXC.1.1        CATAAATAGGGGCGGAGCCGCCTCCCGCTCTCCAAGGTCCTGAGCGGCTGGGTCCCGGCC 480
Hereford_DUXC.1.1      CATAAATAGGGGCGGAGCCGCCTCCCGCTCTCCAAGGTCCTGAGCGGCTGGGTCCCGGCC 480
Holstein_DUXC.1.1      CATAAATAGGGGCGGAGCCGCCTCCCGCTCTCCAAGGTCCTGAGCGGCTGGGTCCCGGCC 480
Tibetan_DUXC.1.1       CATAAATAGGGGCGGAGCCGCCTCCCGCTCTCCAAGGTCCTGAGCGGCTGGGTCCCGGCC 480
Tuli_DUXC.1.1          CATAAATAGGGGCGGAGCCGCCTCCCGCTCTCCAAGGTCCTGAGCGGCTGGGTCCCGGCC 480
Wagyu_DUXC.1.1         CATAAATAGGGGCGGAGCCGCCTCCCGCTCTCCAAGGTCCTGAGCGGCTGGGTCCCGGCC 480
Jersey_DUXC.1.1        CATAAATAGGGGCGGAGCCGCCTCCCGCTCTCCAAAGTCCTGAGCGGCTGGGTCCTGGCC 480
                       *********************************** ******************* ****

Hanwoo_DUXC.1.1        CACTAGGCCAGCAGTCTGCAGCGTCCGTGCGGCTCTCCCACCATGGCTTCGTCCGGCTCC 540
Hereford_DUXC.1.1      CACTAGGCCAGCAGTCTGCAGCGTCCGTGCGGCTCTCCCACCATGGCTTCGTCCGGCTCC 540
Holstein_DUXC.1.1      CACTAGGCCAGCAGTCTGCAGCGTCCGTGCGGCTCTCCCACCATGGCTTCGTCCGGCTCC 540
Tibetan_DUXC.1.1       CACTAGGCCAGCAGTCTGCAGCGTCCGTGCGGCTCTCCCACCATGGCTTCGTCCGGCTCC 540
Tuli_DUXC.1.1          CACTAGGCCAGCAGTCTGCAGCGTCCGTGCGGCTCTCCCACCATGGCTTCGTCCGGCTCC 540
Wagyu_DUXC.1.1         CACTAGGCCAGCAGTCTGCAGCGTCCGTGCGGCTCTCCCACCATGGCTTCGTCCGGCTCC 540
Jersey_DUXC.1.1        CACTAGGCCAGCAGTCTGCAGCGTCCGTGCGGCTCTCCCACCATGGCTTCGTCCGGCTCC 540
                       ************************************************************

Hanwoo_DUXC.1.1        TCCAGCACCTCACGCG 556
Hereford_DUXC.1.1      TCCAGCACCTCACGCG 556
Holstein_DUXC.1.1      TCCAGCACCTCACGCG 556
Tibetan_DUXC.1.1       TCCAGCACCTCACGCG 556
Tuli_DUXC.1.1          TCCAGCACCTCACGCG 556
Wagyu_DUXC.1.1         TCCAGCACCTCACGCG 556
Jersey_DUXC.1.1        TCCAGCACCTCACGCT 556
                       ***************
```

**Supplementary Figure 9.** Clustal Omega multiple sequence alignment of *DUXC* exon 1 across cattle breeds.
